# Tumor-associated tissue-resident macrophages drive pancreatic cancer progression through IGF1-IGF1R signaling

**DOI:** 10.64898/2026.05.08.723679

**Authors:** Yuya Yamamoto, Kenta Takeuchi, Shunsuke Tabe, Ayumu Okumura, Kenji Aoshima, Ryotaro Eto, Takanori Konishi, Naoto Yamamoto, Yohei Miyagi, Masayuki Ohtsuka, Naoki Tanimizu, Hideki Taniguchi

## Abstract

The specific contribution of tissue-resident macrophages (TRMs) to pancreatic ductal adenocarcinoma (PDAC) progression remains unclear. Here, we found that a high abundance of TRM-derived tumor-associated macrophages (TRM-TAMs) is an independent indicator of poor prognosis in patients with PDAC. To elucidate the underlying mechanism, we established an advanced organoid platform (iMac-FPCO), which incorporates macrophages derived from human induced pluripotent stem cells to reflect the differentiation process of TRMs. Single-cell transcriptomic analysis revealed this model recapitulates the transcriptional identity of TRM-TAMs in patient tissue. We demonstrated that TRM-TAMs drive cancer cell proliferation, while maintaining chemoresistance, and identified TRM-derived insulin-like growth factor 1 (IGF1) as the critical mediator. This result provides a rationale for why previous trials targeting IGF1 receptor (IGF1R) failed to improve survival in unselected patient populations. We hypothesize that stratifying patients by TRM-TAM abundance could help identify a responsive subgroup, thereby reviving IGF1R-targeted therapy as a viable treatment for PDAC.

## Introduction

Pancreatic ductal adenocarcinoma (PDAC) remains one of the most lethal malignancies worldwide, with a 5-year survival rate of about 13% despite advances in multimodal therapy.^1, 2^ This dismal prognosis is attributed mainly to its diagnosis at an advanced stage because of limited symptoms, early metastatic dissemination, and resistance to chemotherapy.^3^ A defining characteristic of PDAC is its complex tumor microenvironment (TME), which comprises cancer cells embedded within a dense desmoplastic stroma containing cancer-associated fibroblasts (CAFs), tumor endothelial cells, and diverse types of immune cells.^4, 5^ Among the immune cells, tumor-associated macrophages (TAMs) represent the most abundant immune population^6^ and play multifaceted roles in promoting tumor progression, including angiogenesis, immune suppression, and drug resistance^7, 8^, however, the underlying mechanisms remain unclear.

TAMs originate from two distinct macrophage populations: tissue-resident macrophages (TRMs) derived from erythroid-myeloid progenitors (EMPs) that seed tissues during development and bone marrow-derived macrophages (BMDMs) recruited from the circulation during inflammation and tumor progression.^9, 10^ Traditionally, TAMs are presumed to originate predominantly from BMDMs during inflammation, which often precedes tumor formation, and recruits BMDMs to injured tissues.^11^ However, advances in single-cell RNA sequencing (scRNA-seq) analysis have revealed that TRM-derived TAMs (TRM-TAMs) constitute a significant proportion of the TAM population across various cancers.^12, 13^ The functional roles of TRM-TAMs in the TME remain controversial, as they can act as tumor promoters or suppressors, depending on the tumor type.^14^ In breast cancer, TRM-TAMs induce a cytotoxic T cell response and exert tumor-suppressive effects.^15^ In contrast, in non-small cell lung cancer, TRM-TAMs are involved in the early stages of tumor invasion and act as tumor-promoting factors by inducing regulatory T cells.^12^ In PDAC, TRM-TAMs can be identified by the specific marker genes *FOLR2* and *LYVE1*.^16^ They play an important role in shaping the TME of PDAC. TRMs form a pro-tumorigenic niche during early carcinogenic stages,^17^ contribute to desmoplasia through collagen deposition through the activation of pancreatic stellate cells,^18^ and mediate chemotherapy resistance via metabolic reprogramming.^19^ These studies indicate the pro-tumorigenic functions of TRM-TAMs, they have been primarily conducted in mouse models. However, this leaves a critical translational gap, as the precise molecular mechanisms driving these effects in the human TME remain largely uncharacterized.

A major obstacle in understanding the specific contributions of TRM-TAMs in PDAC is the absence of valid model systems. Conventional co-culture models have used blood-derived macrophages or monocytic cell lines, such as THP-1,^20^ which cannot recapitulate the unique properties of embryonic-derived TRMs. While our previous work incorporating THP-1-derived macrophages into an organoid platform revealed essential roles for monocyte-derived macrophages in angiogenesis and cancer cell survival in a malnourished environment, ^21^ this approach could not address TRM-specific functions. Alternatively, isolating primary human TRMs for ex vivo culture, also presents significant challenges, as these cells rapidly lose their characteristic properties when removed from their native tissue environment, which prevents a detailed functional analysis.^22^ These findings highlight a need for a model that provides both a physiologically relevant cell source and a supportive 3D microenvironment.

To address this challenge, we generated TRM-like cells from human induced pluripotent stem cells (hiPSCs) through EMPs and then integrated these hiPSC-derived macrophages (iMacs) into our established multicellular organoid platform.^24, 25^ This novel system was designed iMac-FPCO. This physiologically relevant model enabled us to identify the mechanism underlying our clinical finding that TRM-TAM abundance is correlated with poor prognosis. We found that these TRM-TAMs actively promote cancer cell proliferation through IGF1-IGF1R signaling. These results provide new mechanistic insights into the pro-tumorigenic functions of TRM-TAMs in PDAC and suggest strategies for therapeutic interventions.

## Results

### High accumulation of TRM-TAMs is associated with poor prognosis in PDAC patients

Previous studies, including ours, have demonstrated that macrophages are abundantly infiltrated in PDAC tissue.^6, 21^ Recent scRNA-seq data suggest that TAMs derived from both BMDMs and TRMs are present in PDAC tissue.^16, 18^ However, the clinical significance of these distinct origins remains poorly defined. To address this, we first performed immunofluorescence staining for CK19 (cancer cell), CD68 (pan-macrophage), and FOLR2 (TRM marker) on tissue samples from 74 PDAC patients who underwent surgical resection between 2015 and 2018. This initial histological analysis revealed marked heterogeneity in the accumulation of FOLR2^+^ TRM-TAMs among the samples, with some tumors exhibiting dense infiltration while others showed only sparse populations (Figure 1A). To systematically analyze this observation and determine its correlation with clinical outcomes, we defined a “TRM ratio” as the percentage of FOLR2^+^CD68^+^ cells among all CD68^+^ macrophages within the tumor areas. To establish a clinically relevant cutoff for the TRM ratio, we first examined its distribution in our patient cohort. The histogram revealed a non-normal, bimodal-like distribution, suggesting distinct populations separated by a natural valley near the 10% mark (Figure S1A). Stratification was supported by a public scRNA-seq dataset, which indicated the presence of a population with a high TRM ratio (Figure S1B). To determine the optimal threshold using statistical validation, a receiver operating characteristic (ROC) curve analysis was performed to predict 1-year survival. This analysis confirmed that a cutoff of 9.6% was optimal (AUC = 0.69, p = 0.031), and it provided statistical support for our stratification method (Figure S1C). Based on these combined findings, we used a cutoff value of 10% to stratify patients into high and low TRM groups. This resulted in the classification of 33 patients (45%) into the high-TRM group and 41 patients (55%) into the low-TRM group. Univariate analysis confirmed that at the time of surgery, baseline clinicopathological characteristics such as age, sex, and tumor stage were well-matched between the two groups (Table 1). There was no significant difference in the total number of CD68^+^ macrophages between the two groups (p=0.37), indicating that the classification was based on the proportion of TRMs in the tumor tissue rather than on overall macrophage infiltration (Figure 1B). Survival analysis revealed significant differences in the clinical outcomes of the two groups. Kaplan–Meier curves demonstrated that patients with the high TRM group exhibited significantly shorter overall survival (OS; median: 22.2 months, 95% confidence interval (CI): 14.3–30.3) compared to the low-TRM group (median: not reached, 95% CI: 24.0–not reached; p = 0.00094, Figure 1C). Similarly, the high TRM group exhibited significantly shorter disease-free survival than the low TRM group (p=0.0030, Figure 1C). In addition, post-operative surveillance revealed that peritoneal dissemination recurrence was significantly more frequent in the high TRM group (p=0.018, Table 1). To determine whether the TRM ratio represents an independent prognostic factor, a multivariate Cox regression analysis was performed by adjusting for the established prognostic factors. After adjusting for pathological stage and histological grade, a high TRM ratio remained an independent predictor of poor OS (HR = 2.76, 95% CI: 1.41–5.40, p = 0.003; Figure 1D). This strong clinical association raises the critical question of how TRM-TAMs mechanistically contribute to a poor prognosis. However, mechanistically interrogating these functions is not feasible using patient samples alone. To determine the functional role of TRM-TAMs in driving this aggressive phenotype, a tractable experimental system that can recapitulate the human TME is essential.

**Figure 1.**
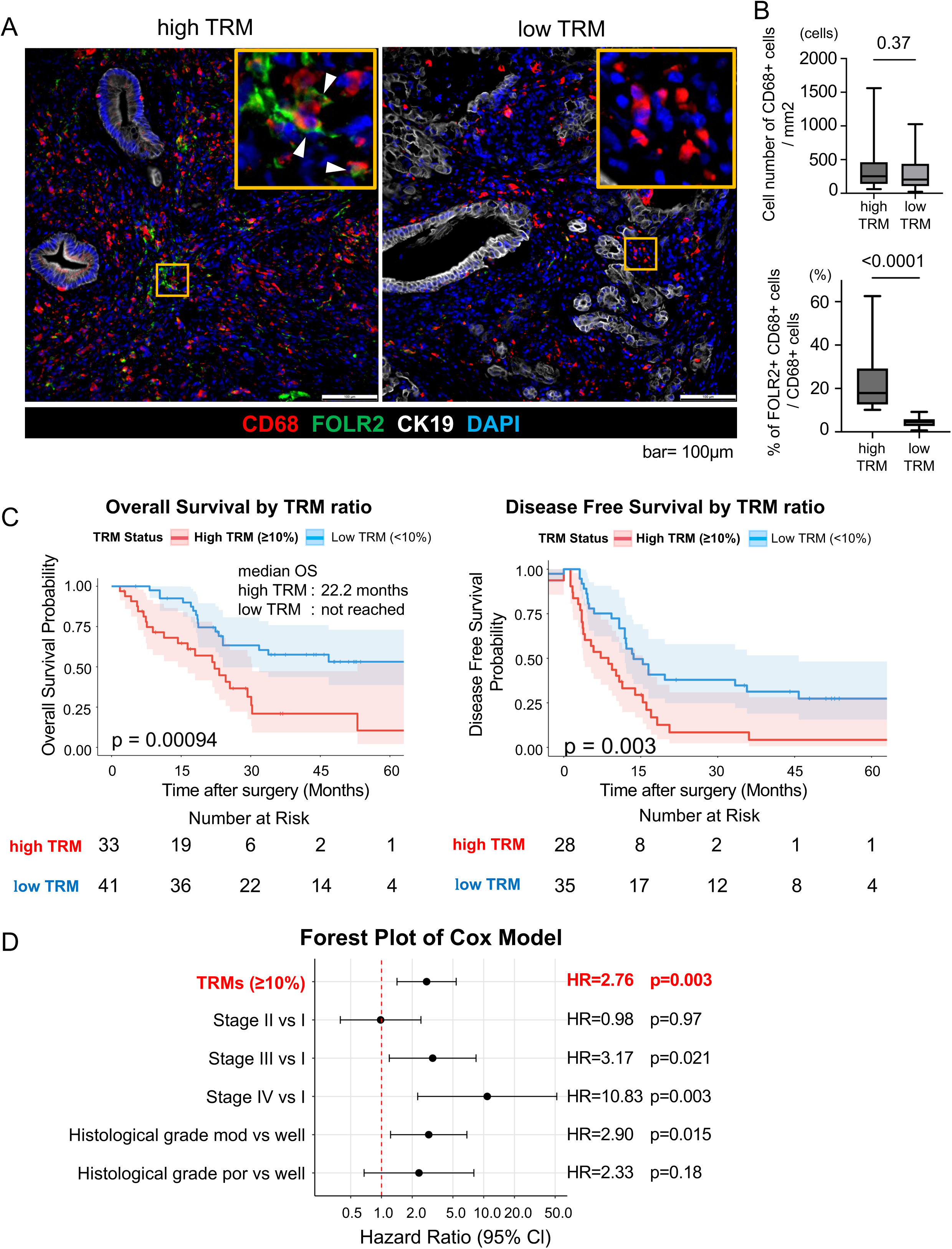
High accumulation of TRM-TAMs is associated with poor prognosis in PDAC patients. (A) Representative immunofluorescence images of high TRM (top) and low TRM (bottom) patient tumors. The stains included CK19 (cancer cells, white), CD68 (pan-macrophages, red), FOLR2 (TRMs, green), and DAPI (nuclei, blue). Arrowheads indicate FOLR2^+^CD68^+^ cells. Scale bar, 100 µm. (B) Quantification of total CD68^+^ macrophages (top) and the TRM ratio (bottom) in high (n=33) and low (n=41) TRM patient groups. Box plots show the median, interquartile range, and min/max values. (C) Kaplan-Meier analyses of OS (left) and DFS (right) for high TRM vs. low TRM patient groups. Median survival times and the number of patients at risk at each time point are indicated. (D) Forest plot of a multivariate Cox proportional hazards model for OS. The analysis was adjusted for established prognostic factors, including pathological stage and histological grade. HR and 95% CI are shown for each variable. Statistical significance was determined by the Mann–Whitney U test (B) or log-rank test (C). See also Figure S1. CI, confidence interval; HR, hazard ratio; OS, overall survival; TAM, tumor-associated macrophage; TRM, tissue-resident macrophage; TRM-TAM, TRM-derived TAM.

### Establishment of a PDAC organoid model incorporating hiPSC-derived macrophages

To create a tractable platform for examining the function of TRM-TAMs in the PDAC TME, we used our fused pancreatic cancer organoid (FPCO) system^24, 25^ which recapitulates PDAC tissue with the TME. As a source of TRMs, we induced macrophages from hiPSCs via EMPs, which are the origin of TRMs^26^ (Figure 2A). iMacs were differentiated using a 21-day protocol, and CD14^+^CD45^+^ macrophages were acquired at high purity (96.8%, Figures 2B and S2A). The iMacs were co-cultured with patient-derived pancreatic cancer organoids (PCOs), hiPSC-derived endothelial cells (ECs), and mesenchymal cells (MCs) in ultra-low attachment plates. They were then transferred to an air–liquid interface environment to produce FPCO containing iMacs (iMac-FPCO, Figure 2C).

**Figure 2.**
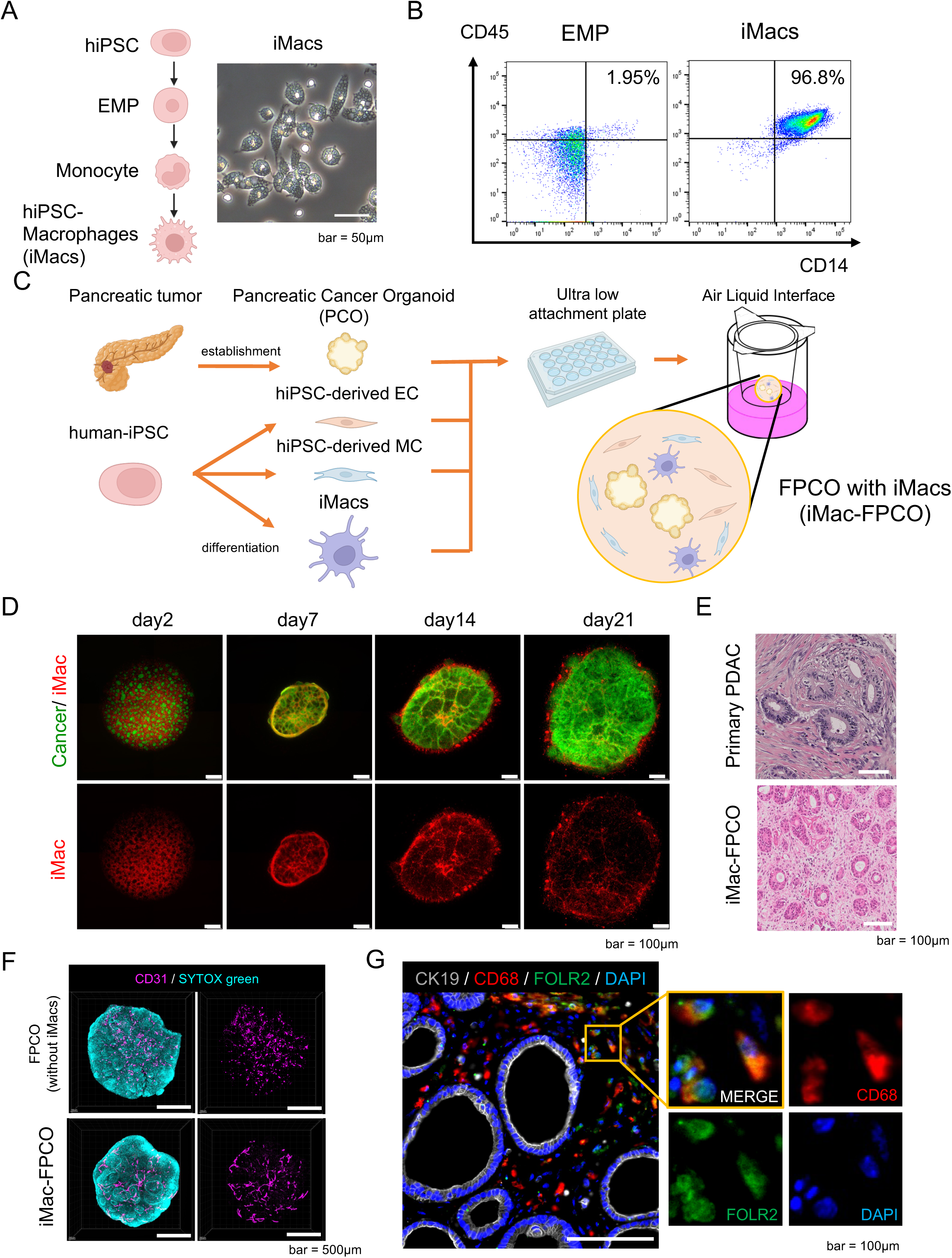
Establishment of a PDAC organoid model incorporating hiPSC-derived macrophages. (A) Schematic of iMacs differentiation (left) and a brightfield image of mature iMacs (right). Scale bar, 25 µm. (B) Flow cytometry analysis showing the purity of iMacs on Day 21 of differentiation, with high expression of macrophage markers CD14 and CD45 (96.8%). (C) Schematic of iMac-FPCO generation, co-culturing of a patient cancer organoid with iMacs and hiPSC-derived stromal cells (ECs and MCs) and maintained at the air–liquid interface. (D) Time-course imaging of iMac-FPCO organization. Cancer cells (GFP, green), iMacs (Kusabira Orange, red). Scale bar, 100 µm. (E) H&E staining comparing the histology of iMac-FPCO with primary human PDAC tissue. Scale bar, 100 µm. (F) Whole-mount immunofluorescence of the iMac-FPCO and FPCO (without macrophages), showing the complex 3D architecture, including CD31^+^ vascular-like networks (magenta). Scale bar, 500 µm. (G) Immunofluorescence staining confirming stable integration of macrophages (CD68, red) that maintain the expression of the TRM marker FOLR2 (green). Scale bar, 100 µm. See also Figure S2. EC, endothelial cell; EMP, erythromyeloid progenitor; H&E, hematoxylin and eosin; hiPSC, human induced pluripotent stem cell; FPCO, fused pancreatic cancer organoid; iMac-FPCO, FPCO with iMacs; iMac, hiPSC-derived macrophage; MC, mesenchymal cell; PDAC, pancreatic ductal adenocarcinoma; TRM, tissue-resident macrophage.

The resulting iMac-FPCO was successfully maintained for up to 21 days, with progressive structural organization (Figure 2D). Macrophages remained integrated throughout the culture period along with cancer cells. Histological analysis revealed that the iMac-FPCO recapitulated the architecture of primary PDAC tissue, with dense stromal compartments and ductal structures (Figure 2E). Whole-mount imaging revealed a complex three-dimensional architecture including vascular-like structures that were positive for CD31 (Figure 2F). Immunofluorescence staining revealed CD68^+^ macrophages throughout the organoid, which maintained the expression of FOLR2, a typical marker of the TRM phenotype (Figure 2G). These results indicate that iMac-FPCO recapitulates the key features of the PDAC TME, including FOLR2^+^ TRM-TAMs and an organized tissue architecture suitable for studying TRM-TAM function during the progression of PDAC.

### scRNA-seq reveals that macrophages in iMac-FPCO recapitulate patient TRMs

Before using this model for functional studies, it was important to systematically validate that the macrophages within the iMac-FPCO accurately recapitulate the transcriptional identity of the patient TRM-TAMs. We performed a multi-step comparative analysis (Figure 3A). First, we performed scRNA-seq on the iMac-FPCO on Day 7 and Day 14, identifying the following four major cell populations: cancer cells, CAFs, TAMs, and endothelial cells (Figures 3B and S3A). As a reference, we used a PDAC-TAM atlas derived from a large patient dataset^27^ by applying previously published gene sets^16^, which classified TAMs into seven distinct clusters including PDAC-FOLR2 TRM-TAM (Figures 3C, S3B and S3C). The PDAC-C3 TAM cluster could not be classified by signature score but exhibited a signature score pattern similar to PDAC-FOLR2 TRM-TAM (Figure S3B). This cluster was considered to be a population related to the previously reported TRM-TAM expressing high levels of *C3* and *CX3CR1* ^28^.

**Figure 3.**
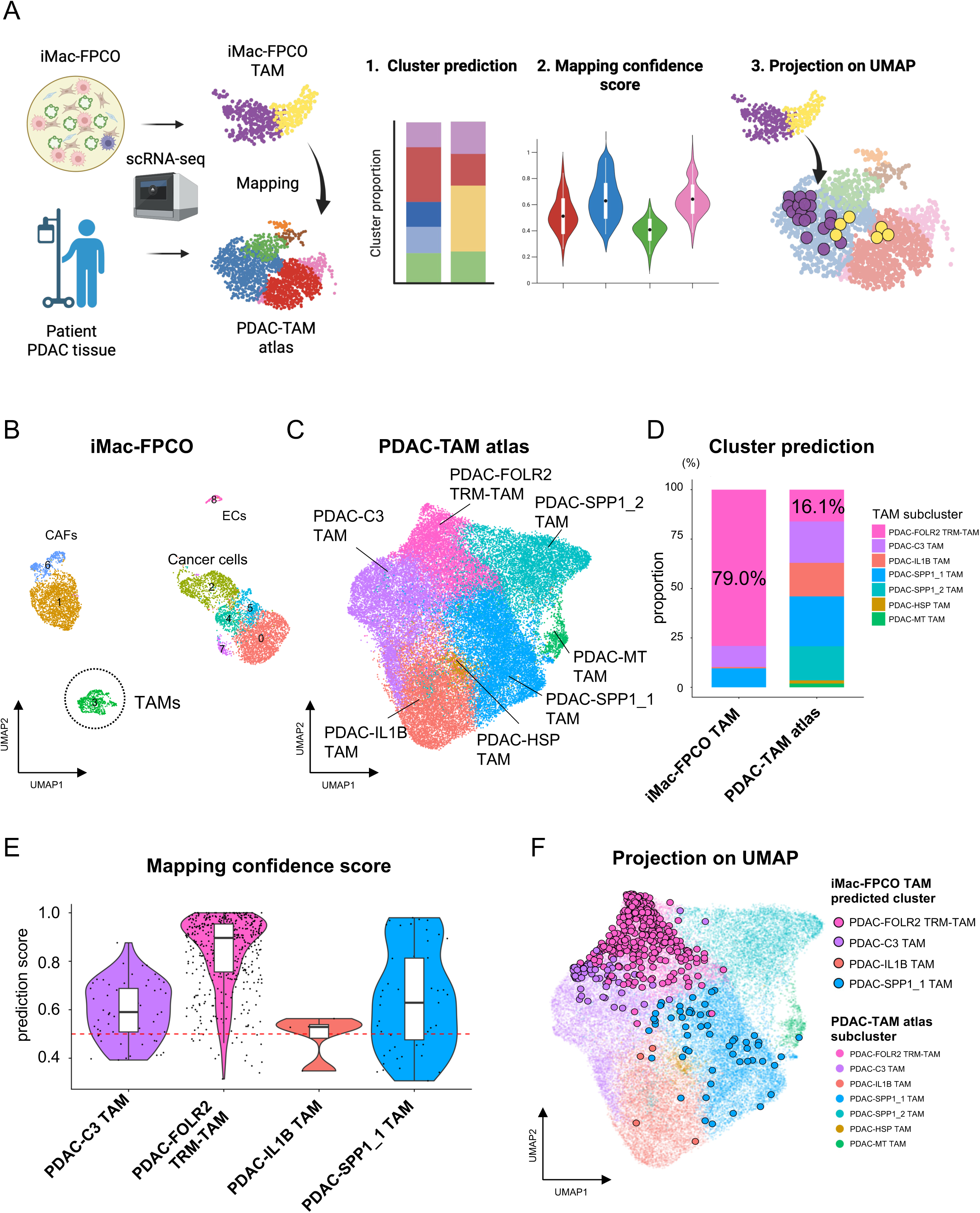
scRNA-seq reveals that macrophages in iMac-FPCO recapitulate patient TRMs. (A) Schematic of the computational workflow for comparing iMac-FPCO TAMs with PDAC-TAM atlas. (B) UMAP plot of iMac-FPCO scRNA-seq data showing the major cell populations. (C) UMAP plot of the reference patient PDAC-TAM atlas, showing seven TAM subclusters. (D) Bar chart showing that the proportion of predicted clusters in iMac-FPCO TAM and the original cluster in the PDAC-TAM atlas (E) Violin plot showing the distribution of mapping confidence scores for each predicted cluster in iMac-FPCO. (F) UMAP visualization of iMac-FPCO TAMs colored with predicted clusters onto the PDAC-TAM atlas UMAP. See also Figure S3. scRNA-seq, single cell RNA sequencing; FPCO, fused pancreatic cancer organoid; iMac-FPCO, FPCO with iMacs; hiPSC, human induced pluripotent stem cell; iMac, hiPSC-derived macrophage; TAM, tumor-associated macrophage; PDAC, pancreatic ductal adenocarcinoma; UMAP, Uniform Manifold Approximation and Projection.

To verify whether iMac-FPCO TAMs recapitulate TRM-TAMs in the PDAC-TAM atlas, we adopted a robust reference-based mapping strategy.^29^ This involved assigning iMac-FPCO TAMs to a predicted cell identity and projecting them onto the established PDAC-TAM atlas Uniform Manifold Approximation and Projection (UMAP) space. First, we determined whether iMac-FPCO TAMs show a general transcriptional similarity to the in vivo PDAC-TAM population as a whole. We examined the overall distribution of mapping confidence scores and found that they were exceptionally high, with most cells scoring above 0.75 with a pronounced peak near 1.0 (Figure S3D). This finding indicates a global resemblance between the iMac-FPCO TAMs and the in vivo patient cell states, thus validating the robustness of the mapping analysis approach. Next, we determined which specific patient subtype closely approximated the iMac-FPCO cells. The cell composition of the iMac-FPCO (predicted proportions) and the PDAC-TAM atlas (original composition) revealed that the majority (79.0%) of iMac-FPCO TAMs were computationally predicted to the PDAC-FOLR2 TRM-TAM cluster (Figure 3D). To validate the specificity of this mapping result, we evaluated the quality of the prediction by examining the confidence scores for each predicted cluster. The cells mapped to PDAC-FOLR2 TRM-TAM showed a median mapping confidence score of nearly 1.0, indicating similar transcriptional identity (Figure 3E). In contrast, the few cells that mapped to other clusters, such as PDAC-SPP1_1 TAM, exhibited widely dispersed and lower confidence scores (median score < 0.6). This finding indicates that mapping to PDAC-FOLR2 TRM-TAM is not merely a best-guess assignment by the algorithm but represents a true transcriptional approximation. Finally, to visually substantiate these conclusions, we projected the iMac-FPCO TAMs onto the PDAC-TAM atlas UMAP coordinates. The iMac-FPCO TAMs co-localized precisely within the spatial coordinates defined by the patient PDAC-FOLR2 TRM-TAM cluster (Figure 3F). This finding indicates that the iMac-FPCO TAMs and the patient-derived PDAC-FOLR2 TRM-TAM population occupy the same location in the reference-defined transcriptional space, confirming that the model is a specific and high-fidelity representation of PDAC patient-derived TRM-TAMs.

To further validate the unique TRM-like identity of the TAMs in iMac-FPCO, we compared them with TAMs from the M0-FPCO, which is our previously established organoid model incorporating BMDM-like THP-1 cells^20^. Integrated UMAP plot showed that TAMs from iMac-FPCO and M0-FPCO formed distinct clusters (Figure S3E). To characterize the identity of these distinct populations, we scored each using an FOLR2-TAM gene signature. The analysis revealed that TAMs in iMac-FPCO showed a higher FOLR2-TAM signature score compared with those in the M0-FPCO (Figure S3F). This finding demonstrates that the source of macrophages (embryonic-like iMacs versus monocytic THP-1 cells) determines the resulting TAM phenotype, and confirms that the iMac-FPCO platform successfully recapitulates a TRM-TAM-like identity, which contrasts with models based on bone marrow derived monocytic precursors.

Taken together, these multifaceted analyses indicate robust evidence that our iMac-FPCO model successfully recapitulates the transcriptional identity and key subpopulations of patient-derived TRM-TAMs, thus establishing it as a high-fidelity platform for investigating their function.

### Trajectory analysis reveals a branched differentiation of TRMs in iMac-FPCO

A key advantage of our FPCO model is its ability to maintain PDAC tissue associated with TME over the long term, because of the endogenous, continuous supply of ECM and growth factors. This feature enables us to track cellular changes over time. Therefore, to understand how TRMs exert their function, we evaluated the heterogeneity and developmental relationships among the validated TRM-TAM subpopulations in iMac-FPCO. First, we analyzed the macrophage compartment. Unsupervised clustering classified macrophages into four distinct subpopulations (Figure 4A). To assign biological identities to these clusters, they were scored them using the same gene lists^16^ previously used to construct the PDAC-TAM atlas. Each cluster was subsequently named as follows based on its high-scoring signature: FOLR2 TRM-TAM 1, FOLR2 TRM-TAM 2, IL1β TAM, and proliferating TAM (Figures 4B, 4C, and S4A). This data-driven annotation was further supported by the distinct marker gene profiles for each subpopulation: The FOLR2 TRM-TAM 1 cluster was characterized by high expression of antigen presentation genes (*IFI30* and *HLA-DPB1*), whereas the FOLR2 TRM-TAM 2 cluster expressed mature TRM markers (*LYVE1*, *F13A1,* and *SIGLEC1*; Figures 4D and S4B). The IL1β TAM cluster showed high expression of inflammatory genes (*IL1B*, *CXCL8*, and*CCL2*), and the proliferating TAM cluster expressed cell cycle-related genes (*TOP2A and MKI67*, Figure 4D). Gene Ontology analysis revealed distinct functional profiles: The FOLR2 TRM-TAM 1 cluster was enriched for antigen presentation pathways, the FOLR2 TRM-TAM 2 cluster for tissue homeostasis and remodeling, the IL1β TAM cluster for inflammatory responses and angiogenesis, and the proliferating TAM for the cell cycle (Figure S4C).

**Figure 4.**
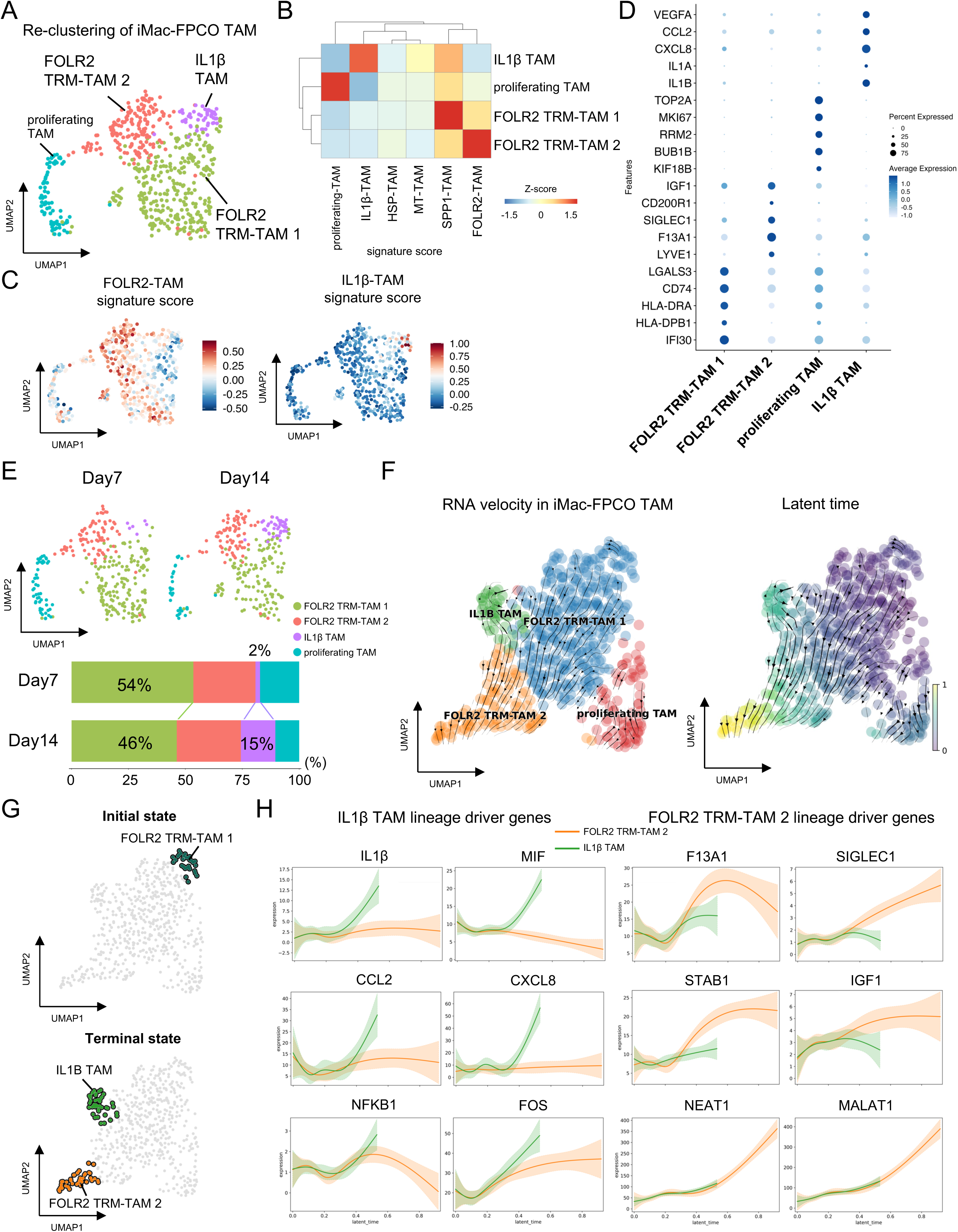
Trajectory analysis reveals a branched differentiation of TRMs in iMac-FPCO. (A) UMAP plot of iMac-FPCO TAMs identifying four distinct subpopulations: FOLR2 TRM-TAM 1, FOLR2 TRM-TAM 2, IL1β TAM, and proliferating TAM. (B) Heatmap showing previously reported signature scores among the four clusters of iMac-FPCO. (C) Feature plots showing enrichment of the FOLR2-TAM and IL1B-TAM signature score in their respective clusters. (D) Dot plot of key marker genes defining each macrophage subpopulation. (E) UMAP plots (top) and bar chart (bottom) showing the distribution of cells from Day 7 (top) and Day 14 (bottom) of culture. (F) RNA velocity analysis projected onto the UMAP (left). Velocity streams (arrows) indicate a branched differentiation trajectory. The UMAP plot colored by latent time (right) provides further support for the proposed developmental trajectory. (G) UMAP plots showing the initial state (FOLR2 TRM-TAM 1, up), and the two terminal states (FOLR2 TRM-TAM 2 and IL1β TAM, bottom) using by CellRank (H) Expression dynamics of IL1β TAM lineage driver genes (left) and FOLR2 TRM-TAM2 lineage driver genes (right) along the inferred latent time axis. See also Figure S4. FPCO, fused pancreatic cancer organoid; iMac-FPCO, FPCO with iMacs; iMac, hiPSC-derived macrophage; hiPSC, human induced pluripotent stem cell; TRM, tissue-resident macrophage; TAM, tumor-associated macrophage.

Having defined these subpopulations, we examined their dynamic transitions in iMac FPCO in culture. Temporal analysis revealed a significant shift in composition between Day 7 and Day 14, with the proportion of IL1β TAM increasing at the expense of the FOLR2 TRM-TAM 1 cluster (Figure 4E). This finding indicates that the macrophage population in iMac-FPCO is not static but evolves dynamically, which suggests that the FOLR2 TRM-TAM 1 cluster represents a progenitor-like state. To formally model the developmental relationships underlying this temporal shift, we performed an RNA velocity analysis^30^. The velocity stream revealed a bifurcated trajectory, with FOLR2 TRM-TAM 1 positioned as a root state from which two distinct differentiation paths emerge: one leading to the mature, homeostatic FOLR2 TRM-TAM 2, and the other to the pro-inflammatory IL1β TAM (Figure 4F). To validate this branching model, we calculated fate probabilities using CellRank^31,32^, which estimates the likelihood of a cell transitioning to a terminal state. The analysis suggested that the FOLR2 TRM-TAM 1 cluster served as the initial state, and the FOLR2 TRM-TAM 2 and IL1β TAM clusters served as the two terminal states (Figure 4G). This result indicates that the FOLR2 TRM-TAM 1 cluster possessed high fate probabilities for both the FOLR2 TRM-TAM 2 and the IL1β TAM lineages, firmly establishing it as a common progenitor population for these two divergent fates. To gain mechanistic insights into this fate decision, we identified lineage driver genes whose expression changed alongside the CellRank-inferred trajectories. Differentiation towards the mature FOLR2 TRM-TAM 2 state was characterized by the upregulation of TRM-related genes, including *F13A1*, *SIGLEC1*, and *STAB1*. In contrast, commitment to the IL1β TAM lineage was associated with the expression of key inflammatory chemokines, such as *IL1B* and *CXCL8*, a fate consistent with the identification of canonical inflammatory transcription factors, including *NFKB1* and *FOS*, as key lineage drivers (Figure 4H, Table 3). Taken together, these analyses reveal a sophisticated, branched differentiation pathway within the iMac-FPCO model, with progenitor-like TRMs diverging into two functionally distinct subsets, one specialized for tissue homeostasis and the other for promoting inflammation.

### TRM-TAMs promote cancer cell proliferation while maintaining chemoresistance

Having established and thoroughly characterized a high-fidelity organoid model that captures TRM-TAM differentiation, we determined the functional effects of these macrophages on cancer cell behavior. Initial long-term culture revealed a notable increase in the overall size of iMac-FPCOs compared to their macrophage-free counterparts (Figure 5A). To systematically quantify this pro-proliferative effect, we compared cancer cell growth between iMac-FPCO and macrophage-free FPCO using a luciferase-based assay (Figure 5B). Luciferase was introduced into PDAC cells in our organoids, which enabled a quantitative estimation of the change in cell number. Significantly enhanced cancer cell proliferation was observed in iMac-FPCO compared to FPCO (Patient 1: 3.0-fold increase at Day 21, p<0.0001; Figures 5C, S5A and S5B). Consistently, immunofluorescence staining for the proliferation marker Ki-67 showed that on Day 21 (Figure 5D), the proportion of Ki-67-positive cancer cells was significantly higher in iMac-FPCO compared to FPCO (Figure 5E). These results indicate that TRM-TAMs drive the proliferation of cancer cells within the organoid.

**Figure 5.**
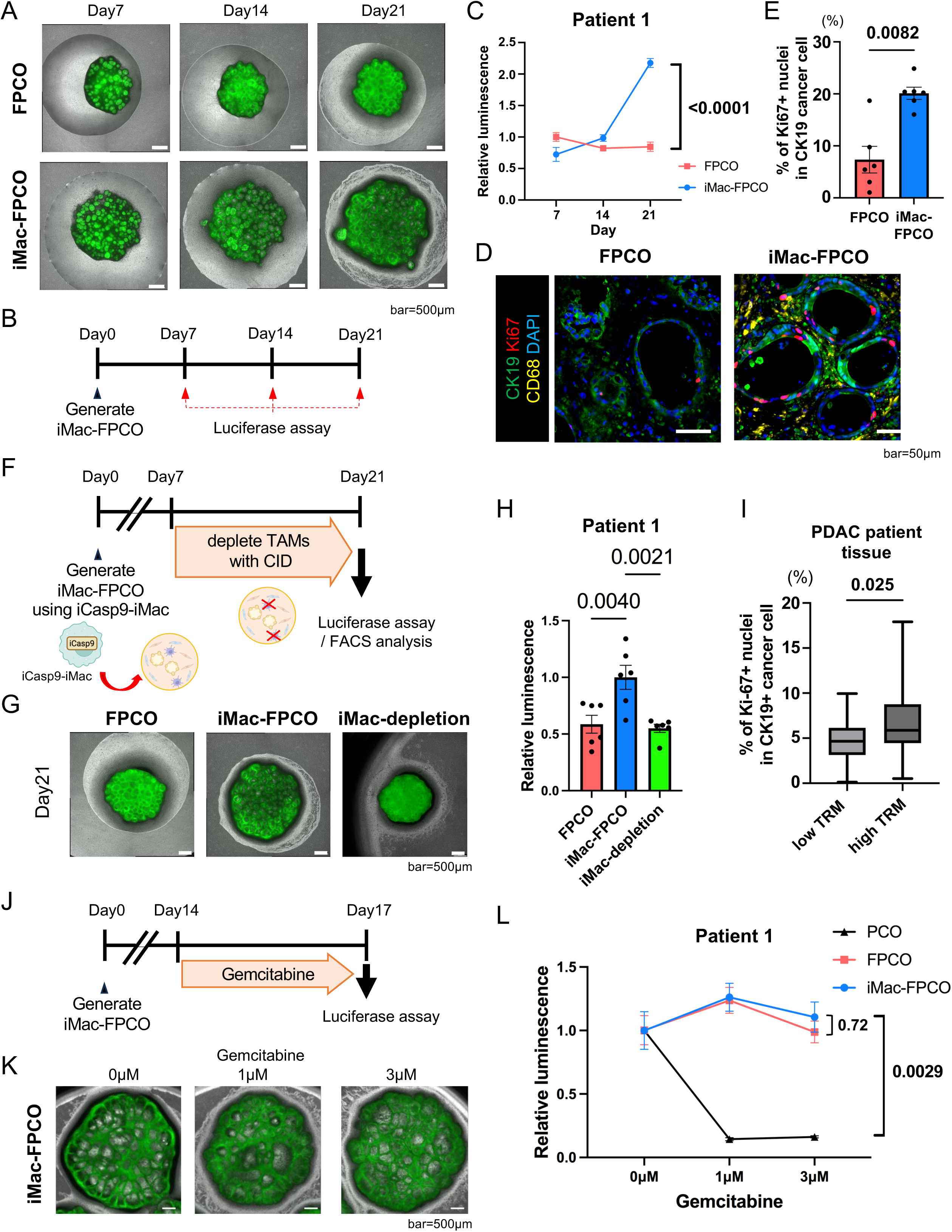
TRM-TAMs promote cancer cell proliferation while maintaining chemoresistance. (A) Representative time-course fluorescence microscope images of FPCO and iMac-FPCO on Day 7, 14 and 21. Cancer cells are labeled with GFP. Scale bar, 500 μm. (B) Schematic of the luciferase-based proliferation assay. (C) Luciferase-based proliferation curves for Patient 1. Organoids were cultured either with iMacs (iMac-FPCO) or without (FPCO) over 21 days. Luminescence values were normalized respectively to the mean of the FPCO and iMac-FPCO group on Day 7. (D) Representative immunofluorescence images of FPCO (left) and iMac-FPCO (right) at Day 21, stained for cancer cells (CK19, green), the proliferation marker Ki-67 (red), and nuclei (DAPI, blue). Scale bar, 50 µm. (E) Quantification of Ki-67 positivity in the CK19^+^ tumor area, confirming higher proliferation in iMac-FPCOs. Data are presented as the mean ± SEM from six biological replicates. (F) Schematic of the iCasp9-based system used for the selective, CID-induced depletion of iMacs within an established iMac-FPCO. (G) Representative fluorescence microscope images of FPCO, iMac-FPCO, and iMac-FPCO treated with CID (iMac-depletion) on Day 21. Cancer cells are labeled with GFP. Scale bar, 500 μm. (H) Quantification of cancer cell proliferation at Day 21 in FPCO, iMac-FPCO and iMac-depletion groups for Patient 1. Luminescence values were normalized to the mean of the iMac-FPCO group at Day 21. (I) Quantification of Ki-67 positivity in the clinical patient cohort, comparing the high TRM (n=31) and low TRM (n=31) groups. The box plot shows the median, interquartile range, and min/max values. (J) Schematic of the gemcitabine resistance assay. Gemcitabine was added to cultures on Day 14. (K) Representative fluorescence microscope images of iMac-FPCO treated with gemcitabine (0, 1 and 3 μM). Cancer cells are labeled with GFP. Scale bar, 500 μm. (L) Quantification of cancer cell viability in response to increasing concentrations of gemcitabine in PCO, FPCO, and iMac-FPCO cultures for Patient 1. Luminescence values from five independent cultures for PCO, FPCO, and iMac-FPCO groups were normalized to the mean of their respective 0 μM control. For all quantitative organoid assays (C, H, and L), the data are presented as the mean ± SEM of 5-6 biological replicates. Statistical significance was determined by a two-way ANOVA with Sidak’s multiple comparisons test (C, L), one-way ANOVA with Dunnett’s multiple comparisons test (H), or the Mann-Whitney U test (E, I). See also Figure S5. iCasp9, inducible Caspase-9; CID, chemical inducer of dimerization; TRM-TAM, TRM-derived TAM; TAM, tumor-associated macrophage; TRM, tissue-resident macrophage; FPCO, fused pancreatic cancer organoid; iMac-FPCO, FPCO with iMacs; iMac, hiPSC-derived macrophage; hiPSC, human induced pluripotent stem cell; GFP, green fluorescent protein; SEM, standard error of mean; ANOVA, analysis of variance.

To confirm the involvement of TRM-TAMs in promoting cancer cell proliferation, we used an inducible cell-depletion system in iMac-FPCO. Macrophages were derived from hiPSCs engineered to express inducible caspase-9 (iCasp9)^33^, which allows for selective depletion upon chemical inducer of dimerization (CID) treatment (Figures 5F and S5C). CID treatment was initiated on Day 7, and the depletion of TRM-TAMs was confirmed by the disappearance of CD45-positive cells with flow cytometry analysis on Day 21 (Figure S5C). Macrophage depletion significantly reduced cancer cell proliferation (Patient 1: 45% reduction, p=0.0021; Figures 5G, 5H, S5E and S5F), which confirmed that TAMs in iMac-FPCO actively promote cancer cell proliferation.

To investigate whether macrophage-driven proliferation observed in our model reflects clinical reality, we re-examined our initial patient cohort. The analysis revealed consistency between our model and patient pathology. Tumors from the high-TRM group showed a significantly higher rate of cancer cell proliferation, as measured by the percentage of Ki-67-positive nuclei, compared to the low-TRM group (p = 0.025, Figures 5I and S5G). This result provides strong clinical evidence that the pro-proliferative function of TRM-TAMs, which we mechanistically established in our organoid system, is a key feature of the aggressive disease phenotype of patients.

Next, we determined whether PDAC cells in iMac-FPCO, which exhibit an increased proliferative capacity, are resistant to drugs such as gemcitabine that target proliferating cells. The administration of gemcitabine commenced on Day 14, when the proliferative activity of the cancer cells became increasingly evident (Figure 5J). Despite increased proliferation, gemcitabine resistance of iMac-FPCO remained comparable to FPCO (Figures 5K, 5L, S5H and S5I), indicating that TAMs within iMac-FPCO promote cancer cell proliferation without sensitizing cancer cells to chemotherapy.

### IGF1-IGF1R signaling mediates TRM-TAM**-**induced cancer cell proliferation

To identify the molecular mechanisms underlying the TRM-TAM mediated proliferation of cancer cells, we conducted a cell-cell interaction analysis using CellChat^34^ on scRNA-seq data from iMac-FPCO and primary PDAC tissue from patients. Our systematic approach involved four steps: extracting cell populations from organoids and patient tissues, predicting cellular interactions, identifying key crosstalk influencing cancer cell proliferation, and screening for inhibitors against the extracted interactions (Figure 6A). CellChat analysis suggested complex interaction networks between the different cell types in both iMac-FPCO and patient samples. We focused on interactions from TAMs and CAFs to cancer cells, identifying numerous potential ligand-receptor pairs (Figures 6B and S6A). Of these, ligands such as *SPP1, IGF1*, and *GRN* were predicted to be highly expressed by the FOLR2 TRM-TAM, while expression of *MIF* and *THBS1* was characteristic of the IL1β TAM (Table 2).

**Figure 6.**
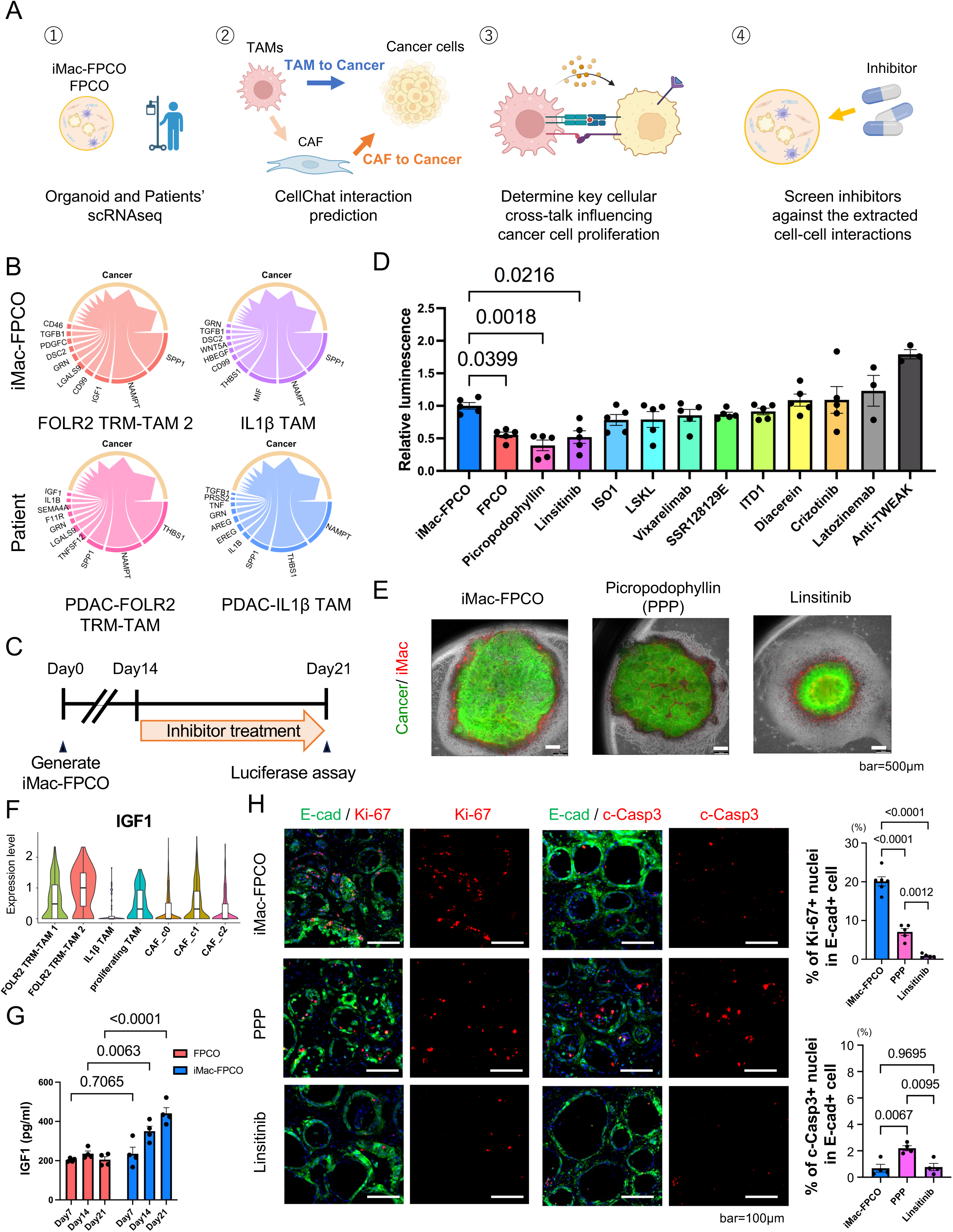
IGF1-IGF1R signaling mediates TRM-induced cancer cell proliferation. (A) Schematic of the systematic approach used to identify key signaling pathways, combining in silico prediction (CellChat) with functional inhibitor screening. (B) CellChat analysis of predicted ligand-receptor interactions from TAM subsets to cancer cells, highlighting conserved pathways between the iMac-FPCO model (top) and patient tissue (bottom). (C) Schematic of the experimental design for the inhibitor screen, where 11 inhibitors targeting predicted pathways were added to iMac-FPCO cultures on Day 14. (D) Bar chart summarizing the results of the inhibitor screen. Cancer cell proliferation was measured on Day 21. Data are presented as the mean ± SEM from 3–5 biological replicates. Luminescence values were normalized to the mean of iMac-FPCO (control). (E) Representative fluorescence microscope images confirming the growth-inhibitory effects of IGF1R inhibitors. Scale bar, 500 μm. (F) Violin plot quantifying IGF1 gene expression in the TAMs and CAFs identified by scRNA-seq in iMac-FPCO. (G) ELISA quantification of secreted IGF-1 protein in conditioned media. Data are presented as the mean ± SEM from four biological replicates. (H) Representative immunofluorescence images showing staining for proliferation (E-cad, green; Ki-67, red; left panels) and apoptosis (E-cad, green; c-Casp3, red; right panels) in iMac-FPCOs treated with vehicle, Linsitinib or PPP. Bar charts show quantification of Ki-67^+^ and c-Casp3^+^ nuclei in E-cad^+^ cancer cells. Data are presented as the mean ± SEM. Scale bar, 100 µm. Statistical significance was using a one-way ANOVA with Dunnett’s multiple comparisons test (D) and (I), one-way ANOVA with Tukey’s multiple comparison test (H), or two-way ANOVA with Sidak’s multiple comparisons test (G). See also Figure S6. FPCO, fused pancreatic cancer organoid; iMac-FPCO, FPCO with iMacs; iMac, hiPSC-derived macrophage; hiPSC, human induced pluripotent stem cell; SEM, standard error of mean; IGF1, insulin-like growth factor 1; CAF, cancer-associated fibroblast; TAM, tumor-associated macrophage; TRM, tissue-resident macrophage; scRNA-seq, single cell RNA sequencing; ANOVA, analysis of variance; c-Casp3, cleaved Caspase 3; E-cad, E-cadherin;, PPP, picropodophyllin.

Based on the predicted interactions and their expression levels, we selected 11 inhibitors targeting key ligand-receptor pairs for functional screening (Table 3). The inhibitors were added to iMac-FPCO on Day 14, and cancer cell proliferation was assessed on Day 21 using a luciferase assay (Figure 6C). Among all tested compounds, IGF1R inhibitors showed the most potent effects. Picropodophyllin (PPP) and Linsitinib, respectively, reduced cancer cell number by 46% (p = 0.0018) and by 42% (p = 0.022), whereas the other inhibitors, including those targeting IL1B, MIF, THBS1, and other pathways showed minimal effects (Figure 6D and Table 3). Representative microscopy images confirmed the growth-inhibitory effects of the IGF1R inhibitors (Figure 6E). Furthermore, IGF1R inhibitors suppressed cancer cell proliferation in different patient-derived organoids (Figures S6B and S6C). In contrast, treatment of FPCO (macrophage-free) with Linsitinib and PPP did not affect cancer cell number (Figure S6D). This finding confirms that inhibitor efficacy is dependent on the presence of TRM-TAMs and strongly indicates that their primary mechanism is the blockade of TRM-TAM derived pro-proliferative signals, not a direct cytotoxic effect of the cancer cells.

To identify the cellular source of IGF1, we examined the expression levels in iMac-FPCO model and patient data. A quantitative analysis confirmed that FOLR2 TRM-TAM was the predominant source of *IGF1* in both iMac-FPCO and patient tissues (Figures 6F and S6E). Validating this at the protein level, ELISA analysis of conditioned media revealed significantly higher IGF1 concentrations in the supernatants from iMac-FPCO compared with FPCO (Figure 6G). Next, to elucidate the mechanism of action underlying the efficacy of IGF1R inhibitors, immunofluorescence staining was performed on iMac-FPCOs treated with Linsitinib and PPP. Consistent with the luciferase assay, Ki-67 quantification revealed a significant reduction in the percentage of Ki-67^+^ cancer cells in both Linsitinib- and PPP-treated organoids compared with the vehicle controls (Figure 6H). In contrast, analysis of the apoptosis marker cleaved caspase 3 (c-Casp3) showed a slight increase in apoptotic cells with PPP treatment, and no significant change with Linsitinib (Figure 6H). Collectively, our systematic approach, combining computational prediction with functional validation, identifies IGF1-IGF1R signaling as an important mediator of TRM-TAM-induced cancer cell proliferation.

## Discussion

In this study, we established a definitive link between TRM-TAMs and clinical outcomes in PDAC, bridging clinical observation with molecular mechanisms (Figure 7). We found that the TRM-TAM accumulation is an independent predictor of poor prognosis. We further dissected the molecular drivers of this phenomenon and demonstrated that TRM-TAMs secrete IGF1, which promotes the PDAC cell proliferation.

**Figure 7.**
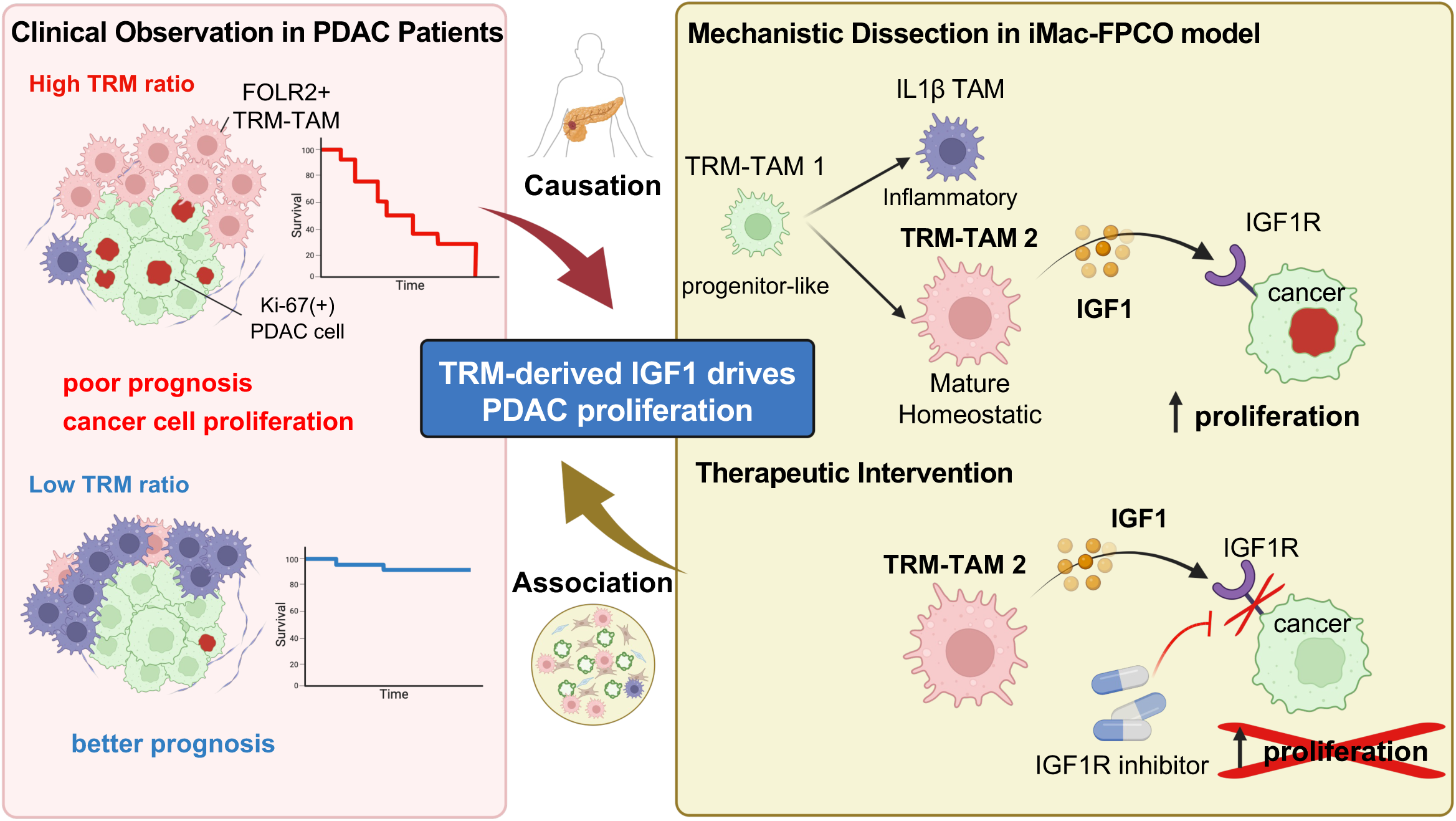
A model for TRM-driven PDAC progression and therapeutic targeting. Schematic model summarizing the study’s findings. Left (Clinical Observation): In PDAC patients, a high TRM ratio is associated with increased cancer cell proliferation (Ki-67^+^) and is an independent predictor of poor clinical prognosis. Top Right (Mechanistic Dissection): The iMac-FPCO model recapitulates the PDAC microenvironment and reveals a developmental trajectory for TRMs. Progenitor-like TRMs (TRM1) mature into a homeostatic, pro-tumorigenic state (TRM2). These mature TRM2 cells are the primary source of secreted IGF1. Bottom Right (Therapeutic Intervention): TRM-derived IGF1 acts on IGF1R expressed by cancer cells, driving their proliferation. This pro-tumorigenic axis may be effectively blocked by small-molecule IGF1R inhibitors. FPCO, fused pancreatic cancer organoid; iMac-FPCO, FPCO with iMacs; iMac, hiPSC-derived macrophage; hiPSC, human induced pluripotent stem cell; PDAC, pancreatic ductal adenocarcinoma; IGF1, insulin-like growth factor 1; TRM, tissue-resident macrophage.

The study of these dynamic cellular processes was facilitated by a significant methodological advance. Our iMac-FPCO platform, which incorporates iMacs, enabled us to elucidate the role of TRM-TAMs in PDAC malignancy, thereby filling a gap left by conventional models. For example, our previous study used the same FPCO platform, but incorporated THP-1 cells as a model for BMDM-derived TAMs.^20^ Although that system successfully recapitulated TAM diversity and identified key contributions to angiogenesis and cancer cell survival in a malnutrition state, it could not address the specific functions of the TRM-TAM population. The current iMac-FPCO model, which incorporates hiPSCs differentiated by the EMP pathway to recapitulate the origin of TRM, provides the first humanized system to specifically isolate this TRM-TAM lineage, which is not adequately represented by conventional monocyte-derived macrophages or cell lines. The physiological relevance of this approach is highlighted by the scRNA-seq analysis, which demonstrated high transcriptional fidelity between TAMs in iMac-FPCO and TRM-TAMs from patient tumors. This high-fidelity system represents a robust platform for mechanistic interrogation, enabling us to identify IGF1-IGF1R signaling as the primary pathway through which TRM-TAMs drive cancer cell proliferation. Taken together, these findings not only deepen our understanding of PDAC biology but also offer a unique window into the complex biology of TRM-TAMs.

The IGF1-IGF1R axis has long been recognized as a promising therapeutic target^35^. While randomized phase II trials initially showed promising results^36^, subsequent phase III studies, such as the GAMMA trial, failed to demonstrate survival benefits in unselected patient populations^37^. This inconsistency highlights a critical limitation in previous strategies: the absence of validated predictive biomarkers. Systemic factors, such as blood IGF1 levels, failed to correlate with clinical outcomes. This left the specific subgroups dependent on this pathway unidentified. Our study offers a new perspective on this clinical problem by identifying TRM-TAMs as a local IGF1-producing niche within the PDAC TME. This result provides a biological rationale for past failures, suggesting that therapeutic response might be dictated by this local TME niche rather than by systemic factors. Therefore, we propose that the TRM ratio, which we established as a robust prognostic factor, may be repurposed as a predictive biomarker to stratify patients and select the subgroup most likely to benefit from IGF1R-targeted therapy. Moreover, our observation that TRM-TAMs fuel proliferation while preserving chemoresistance suggests the potential to combine TRM-TAM-targeted therapies with conventional chemotherapy. This dual effect may be mechanistically explained by the known role of the IGF1-IGF1R axis, in which downstream PI3K/AKT signaling not only to drives proliferation but also confers chemoresistance by providing potent anti-apoptotic survival signals.^38,39^ Thus, TRM-TAMs provide both a “proliferate” and a “survival” signal to cancer cells, a dual function driven by IGF1. Of note, the primary effect of the IGF1R inhibitors, PPP and Linsitinib, was the potent suppression of cancer cell proliferation, rather than the direct induction of widespread apoptosis. The differences observed in the inhibition of cancer cell proliferation may be explained by the distinct pharmacological profiles of the drugs. Linsitinib is a dual inhibitor of both IGF1R and the insulin receptor (INSR)^40^, whereas PPP is more selective for IGF1R.^41^ This dual blockade by Linsitinib may be responsible for its more potent effect on suppressing cancer cell proliferation (Ki-67), as signaling through IGF1R and INSR contributes to mitogenic pathways. In contrast, the modest, yet statistically significant increase in cell death (c-Casp3) observed only with PPP treatment is likely due to its off-target effects, including the induction of mitotic arrest and apoptosis through microtubule depolymerization, which is independent of IGF1R signaling.^42^ Nevertheless, these findings that both inhibitors suppressed TRM-TAM-dependent proliferation indicate that the IGF1R axis is the primary driver of this phenotype.

In addition to providing the correct cell type to dissect the molecular mechanisms underlying PDAC malignancy, the FPCO platform, combined with iMacs, supports the dynamic differentiation into TRM-TAMs *in vitro*. Although previous studies have provided valuable insights into TAM dynamics over time, such as in KPC mouse models,^16^ these have largely been described as mechanisms in murine experimental models. To our knowledge, the temporal dynamics of human TRM maturation and plasticity within the TME have not been fully characterized. Our trajectory analysis, which captured the branched differentiation of progenitor-like TRMs into distinct mature and inflammatory states, highlights the capacity of this to fill this gap and study human TRM biology. This differentiation begins with a common progenitor-like state, FOLR2 TRM-TAM 1, which is characterized by a high antigen presentation capacity, suggesting the potential for initial, although temporary, anti-tumor immune surveillance. From this root, one path leads to the mature state, homeostatic FOLR2 TRM-TAM 2 phenotype, which supports tissue remodeling and homeostasis. The other path leads to a highly pro-inflammatory IL1β TAM phenotype, which promotes inflammation and angiogenesis. This branched model provides a more detailed explanation of TRM plasticity and resolves the previous ambiguity of a linear pathway. Our analysis identified the peak expression of *IGF1* in the FOLR2 TRM-TAM 2 lineage, the subset specializing in tissue homeostasis. This finding suggests that the key pro-proliferative signal for cancer cells is provided not by inflammatory macrophages, but by those adopting a tissue-remodeling and homeostatic role. This result suggests a deceptive form of pro-tumorigenic activity, wherein macrophages engage in seemingly “normal” tissue maintenance but are exploited by the tumor to promote its growth, highlighting a key window and a specific cellular state for therapeutic intervention.

We observed an association between a high TRM ratio and the incidence of peritoneal dissemination recurrence (Table 1). This association might be a direct consequence of the increased tumor burden driven by TRM-mediated proliferation, which is more rapidly growing primary tumor that provides more opportunities for cells to shed and metastasize.^43^ However, it is also possible that TRMs actively facilitate this process through the same “tissue-remodeling” and “homeostatic” programs that we identified. Such functions involve the modulation of the extracellular matrix and creation of a “pre-metastatic niche”.^44^ Therefore, we hypothesize that TRMs may act through a dual mechanism: as local drivers of proliferation that increase metastatic potential and as active facilitators of metastatic colonization. This hypothesis suggests that poor prognosis associated with a high TRM ratio may be multifactorial and reflects a more aggressive tumor beyond IGF1-driven proliferation alone.

The expression of *FOLR2* on pro-tumorigenic TAMs can represent a direct and promising therapeutic target. Future strategies may involve the selective depletion or functional modulation of this cell population.^45^ For instance, one strategy may involve FOLR2-targeted nanoliposomes designed to deliver an IGF1-specific siRNA payload directly to the TRMs.^46^ This approach holds the potential to selectively silence the key pro-tumorigenic factor at its source, effectively disarming the TRM-TAMs without depleting the entire TAM population, while simultaneously overcoming the drug delivery barriers that hinder large-molecule biologics. Moreover, the extensive crosstalk between CAFs and TAMs in the PDAC TME, which is likely important for chemoresistance and immunosuppression, requires further study. Our iMac-FPCO platform, which co-cultures these distinct stromal populations, provides an ideal system to determine how they synergize to create a pro-tumorigenic and treatment-resistant niche, thus offering deeper insights into the biology of this intractable disease. Finally, this iMac-coculturing platform may be able to be applied to other cancers. By co-culturing iMacs with patient-derived organoids from other malignancies, this human-relevant system enables the study of organ-specific imprinting and the context-dependent, often opposing, functions of TRM-TAMs in different cancer types. These future applications will provide further insights into the complex mechanism of cancer biology.

### Limitation of the study

This study had several limitations. First, our retrospective analysis of the TRM ratio in a single-center cohort requires validation. To establish the TRM ratio as a clinically actionable biomarker for patient stratification, future studies must validate its prognostic value in large-scale, multi-center prospective cohorts. Second, the branched differentiation trajectory of TRMs was inferred computationally using RNA velocity analysis. Although this approach provides high-resolution insights into human TRM dynamics that are difficult to capture in vivo, definitive confirmation of these lineage relationships will require the development a tracing system. Third, iMac-FPCO does not contain all the cell types consisting of PDAC TME. We took advantage of this simplified system to intensively analyze cell-cell communication within TME and then identified that TRM-TAMs regulate cancer cell proliferation through the IGF1-IGF1R axis. In future, once T cells can be generated from a patient’s iPSC, it is expecting that FPCOs containing innate as well as adaptive immune cells provide an advanced model in which we may be able to address how CAFs, TAMs, and PDAC cells impair anti-tumor function of T cells in PDAC TME.

## Materials and Methods

### Human tissue samples

Human PDAC patient tissue samples were identified and obtained from Kanagawa Cancer Center and Chiba University. All patients underwent surgery without preoperative chemotherapy. This study was approved by the Institutional Ethics Board of Chiba University Graduate School of Medicine (Ethical approval number #3,302) and Kanagawa Cancer Center (A160128003, 28 疫 55, 2019-6-0716).

### Calculation of the TRM ratio by immunofluorescence analysis and clinical evaluation

TRMs were examined from 74 PDAC patients who underwent surgical resection at Chiba University Hospital from 2015 to 2018. The patients were classified according to the UICC 8th edition TNM staging system. Immunofluorescence staining was done using anti-cytokeratin 19 (MBS423230, 1:200, Mybiosource), anti-CD68 (76437S, Cell Signaling), and anti-FOLR2 (MA5-26933, Invitrogen) antibodies. To ensure objective and reproducible quantification of TRM infiltration, a standardized protocol was established. For each patient sample, 5-15 high-power fields were randomly selected from CK19-positive tumor areas. The analyses ware performed by a single analyst who was blinded to the patients’ clinical outcomes. Quantification was done using a custom-written macro in ImageJ software (v2.14.0) to ensure objectivity and reproducibility. The analytical pipeline was as follows: (1) Multi-channel images were split into individual channels for DAPI (nuclei), CD68 (pan-macrophage), and FOLR2 (TRM). (2) Each channel was converted to a binary mask using a global intensity threshold. (3) The total number of macrophages was determined by counting the particles resulting from the logical intersection (AND operation) of the DAPI and CD68 masks. (4) The number of TRM-TAMs was determined by counting particles from the intersection of the DAPI, CD68, and FOLR2 masks. (5) The TRM ratio for each image was calculated as the number of TRM-TAMs divided by the total number of macrophages, multiplied by 100. The final TRM ratio for each patient represented the average of all analyzed images. To statistically validate this cutoff, a ROC curve analysis was performed for the prediction of 1-year overall survival. The optimal cutoff point was conducted using the Youden’s index.

The associations between TRM ratio and clinicopathological features were analyzed using the Mann-Whitney U test for continuous variables and Fisher’s exact test for categorical variables. Overall survival and disease-free survival were analyzed by the Kaplan-Meier method followed by a Log-rank test. Multivariate Cox proportional hazards regression analysis was used to assess the TRM ratio as an independent prognostic factor, which was adjusted for the UICC 8^th^ stage and histological grade. Statistical significance was defined as p<0.05.

### Calculation of the Ki-67 positivity rate in patient tissue

Fluorescent immunostaining was performed on clinical tissue from 31 patients in the high TRM group and 31 patients in the low TRM group, classified based on the TRM ratio described above. Immunofluorescence staining was performed using anti-cytokeratin 19 (MBS423230, 1:200, Mybiosource), anti-Ki-67 (ab16667, 1:200, Abcam), and anti-FOLR2 (MA5-26933, Invitrogen) antibodies. To ensure objective and reproducible quantification of the Ki-67 positive ratio, a standardized protocol was implemented. For each patient sample, 5-10 high-power fields were randomly selected from CK19-positive tumor areas. All image analysis was performed by a single analyst who was blinded to the TRM ratio classification. Quantification was performed using a custom-written macro in ImageJ software (v2.14.0). The workflow was designed to specifically measure proliferation within cancer cell regions: (1) A cancer cell area mask was generated from the CK19 channel using automated thresholding. (2) Individual nuclei were segmented from the DAPI channel using a combination of Gaussian blur, thresholding, and the watershed algorithm to separate touching nuclei. (3) The total number of cancer cell nuclei (denominator) was determined by counting segmented nuclei that spatially overlapped with the CK19 mask. (4) For each of these nuclei, the mean intensity of the Ki-67 channel was measured. (5) Nuclei with a mean Ki-67 intensity above a predefined, consistent threshold were classified as positive (numerator). (6) The Ki-67 positivity rate was calculated as the number of Ki-67 positive nuclei divided by the total number of cancer cell nuclei, multiplied by 100. Based on classification by TRM ratio, Ki-67 positivity rates were compared using the Mann-Whitney U test.

### Lentivirus preparation

PDAC and hiPS cells were labeled to track each cell population within the organoid. To construct the CSII-EF-Luc-IRES-EGFP plasmid, the firefly luciferase was subcloned into CSII-EF-MCS-EGFP (RIKEN BRC). PDAC cells transfected with CSII-EF-Luc-IRES-EGFP were designated Luc-GFP^+^ PDAC cells. Kusabira-Orange (KuO) was introduced into the hiPS cells, as previously described.^24,25^

### hiPSC-derived macrophage differentiation

hiPSCs were differentiated into macrophages using a 25-day protocol. Briefly, the hiPSCs were initially cultured in AK02 medium (Ajinomoto) (Day -4- 0), followed by StemPro-34 (Thermo Fisher Scientific) medium supplemented with BMP4 (25 ng/ml; R&D systems), VEGF (10 ng/ml; Thermo Fisher Scientific), FGF2 (10 ng/ml; Thermo Fisher Scientific), and CHIR99021 (2 μM; Cayman Chemical Company) for mesoderm induction (Day 0-2). For embryoid body (EB) -like myeloid progenitor (EMP) formation, the cells were cultured in STEMdiff APEL2 (Stem Cell Technologies) containing VEGF (10 ng/ml), FGF2 (10 ng/ml), SCF (50 ng/ml; Peprotech), Flt3L (10 ng/ml; Wako), and TPO (30 ng/ml; Wako) until Day 10. Next, the cells were maintained in serum-free differentiation (SFD) medium^47^ with M-CSF (50 ng/ml; Peprotech) for monocyte differentiation (Day 10-16), followed by StemPro medium with M-CSF (50 ng/ml) for final macrophage maturation on FBS-coated tissue culture plates (Day 16-21). All cultures were maintained at 37°C in 5% CO_2_ with medium changes every 2-3 days. Flow cytometry was used to assess differentiation efficiency using CD45 and CD14 markers, and revealed 96.1% CD45^+^CD14^+^ macrophages by Day 21. Differentiated macrophages were subsequently co-cultured with pancreatic cancer organoids to generate iMac-FPCO.

### PDAC organoid culture

A conventional organoid was established from a PDAC patient as previously described.^24,25^ All patient samples for establishing organoids were acquired from the Kanagawa Cancer Center. The organoid basic medium consisted of DMEM/Ham’s F-12 (Nacalai Tesque, Tokyo, Japan), containing 1% penicillin/streptomycin (Nacalai Tesque, Tokyo, Japan) and GlutaMAX (Thermo Fisher Scientific, Waltham, MA). The PDAC organoid culture medium (complete medium) contained the following factors in the organoid basic medium: B-27 (Thermo Fisher Scientific, Waltham, MA), 10-mM nicotinamide (Sigma-Aldrich, St. Louis, MO), 1-mM N-acetyl-L-cysteine (Sigma-Aldrich, St. Louis, MO), 5 ng/mL of epidermal growth factor (EGF) (Sigma-Aldrich, St. Louis, MO), 0.1 µg/mL of noggin (Pep Rotech, Cranbury, NJ), 0.5-µM A83-01 (Selleck), 0.1 µg/mL of FGF10 (PeproTech, Cranbury, NJ), 10-nM gastrin I (Sigma-Aldrich, St. Louis, MO), 50 ng/mL of Wnt3A (R&D Systems, Minneapolis, MN), and 0.1 μg/mL of R-spondin1 (PeproTech, Cranbury, NJ).

### Establishment of FPCO, including iMacs

The FPCO was established as described previously.^24,25^ iMacs were simultaneously induced for co-culture with patient-derived PDAC cells and hiPSC-derived MCs and ECs. The differentiation protocol from hiPSC to EC and MC was performed as previously described.^24,25^ PDAC cells (1.2 × 10^5^), hiPSC-ECs (8.4 × 10^4^), hiPSC-MCs (2.4 × 10^5^), and hiPSC-Macs (1.2 × 10^5^) were suspended and cultured for 24 h in a 24-well ultra-low attachment plate (Corning Elplasia; Corning) on a basic-FPCO medium, which consisted of a 1:1 mixture of organoid basic medium and endothelial growth medium (EGM, Lonza). After one day, the spheroids were collected and resuspended in a complete-FPCO medium, which consisted of a 1:1 mixture of PDAC organoid medium and EGM. They were dropped onto a cell culture insert (Greiner). The resulting iMac-FPCO was cultured on basic-FPCO medium and maintained for at least 21 days.

### Flow cytometric analysis

hiPS cells, and macrophages were incubated with a FACS buffer containing an Fc block (422,302, BioLegend) for 25 min at 4□. After washing they were resuspended in a FACS buffer containing PE-anti-CD45 antibody (982322, BioLegend) and FITC-anti-CD14 antibody (325608, BioLegend) for 20 min at 4□, followed by 0.1% propidium iodide staining. Expression of CD45 and CD14 was assessed on a BD FACSCelesta^TM^, and the data were processed using FlowJo software (version 10). iMac-FPCO dissociation was achieved by incubation in liver digestion medium^48^ containing 0.5□mg/mL collagenase (Sigma-Aldrich), 0.5□mg/mL pronase (Roche, Basel, Switzerland), and 0.25□mg/mL DNase1 (Sigma-Aldrich) in for 20□min at 37□℃ while shaking. The enzyme activity was terminated by the addition of 5% FBS in PBS. The samples were centrifugted (400 × g) for 5□min to pellet the cells. Next, 100□µl of 5% FBS in PBS was added, and the cells were incubated with PE-anti-CD45 antibody (368509, Biolegend) and SYTOX^TM^ Blue (Invitrogen). Based on the expression of CD45 and GFP, iMacs and Luc-GFP^+^ PDAC cells were detected, respectively.

### Histological analysis and immunofluorescence staining

FPCO and iMac-FPCO were fixed in 4% paraformaldehyde (PFA) and then embedded in paraffin or frozen blocks with 10% sucrose and 7.5% gelatin in phosphate-buffered saline (PBS). Each block was cut into 7-µm-thick sections and stained with hematoxylin and eosin and immunofluorescent markers based on standard histological protocols. The following primary antibodies were used for immunofluorescence staining: anti-cytokeratin 19, anti-CD68, anti-FOLR2, anti-CD31 (M0823, 1:100, Dako), anti-Ki-67 (ab16667, 1:200, Abcam), and anti-Cleaved Caspase-3 (9661, 1:200, CellSignaling). Antigen retrieval was achieved using Tris EDTA buffer (pH 9.0) at 120°C for 12 min. Permeabilization was done using 0.05% Tween20 (Nacalai Tesque) in PBS, and blocking in Protein Block, Serum-Free, Liquid Form (Dako), followed by incubation with primary antibodies at 4°C for 16 h. After washing, secondary antibodies were added and the samples were incubated for 1 h at room temperature. After the secondary antibody reaction, the samples were washed and mounted with Anti-Fade Fluorescence Mounting Medium (ab104135, Abcam) containing 0.1% DAPI (Nacalai Tesque). The following secondary antibodies were used for immunofluorescence staining: donkey anti-goat IgG1 cross-adsorbed secondary antibody, Alexa Fluor™ 488 (Alexa Fluor, A11055); donkey anti-rabbit IgG (H+L) cross-adsorbed secondary antibody, Alexa Fluor™ 555 (Alexa Fluor, A31572); and donkey anti-mouse IgG (H+L) cross-adsorbed secondary antibody, Alexa Fluor™ 647 (Alexa Fluor, A31571). A dilution of 1:400 was used for all secondary antibodies. Images were acquired using a Leica Thunder fluorescence microscope and quantitative analyses were performed using ImageJ (2.14.0) software.

### Whole-mount immunostaining of organoids

The FPCO and iMac-FPCO were fixed in 4 % PFA overnight. After washing, the organoids were delipidated in a 10 % CHAPS solution (Nacalai Tesque) and a 25 % N-methyldiethanolamine (NMDEA) solution (Sigma Aldrich) in a shaking water bath at 37 ◦C for 24 h.^49^ Next, the organoids were washed three times with 0.1 % Tween-PBS (PBST), and the proteins were blocked by incubation with Protein Block, Serum-free (DAKO) for 2 h at room temperature with shaking. The primary antibody anti-CD31 (M0823, 1:100, Dako) was added and incubated for 5 days at 4□ on a rotator. The following day, the organoids were washed with PBST for 24 h. Goat anti-mouse IgG1 cross-adsorbed secondary antibody, Alexa Fluor™ 555 (Alexa Fluor, 1:500, A21127), was added and incubated for 2 days at 4□ on a rotator. The organoids were then washed three times with PBST every 60 min. Sytox green (S34862, 1:1000, Invitrogen) was used to detect the cell nuclei. Dehydration was performed by serial dilution using a methanol solution, and tissue clearing was done using the BABB method as previously described^50^ using a 1:1 mix of benzyl benzoate (Sigma Aldrich) and benzyl alcohol (Tokyo Chemical Industry). Images were captured under a Leica SP8 confocal microscope (Leica). A 3D rendering was generated using Imaris software.

### Enzyme-linked immunosorbent assay (ELISA)

Conditioned media from FPCO and iMac-FPCO cultures were collected on Days 7, 14, and 21. To ensure that the culture medium was free of serum-derived growth factors, the medium was switched to a serum-free basal medium 48 hours before collection. To increase the detection sensitivity of the secreted proteins, the supernatants were concentrated before the assay. Specifically, 600 µL of conditioned media was loaded onto an Amicon® Ultra Centrifugal Filter Unit (Millipore, UFC200324) and centrifuged at 7500× g for 60 min at 4°C. It resulted in an approximate 8-fold concentration of the supernatant. The concentration of IGF-1 in the samples was quantified using a Quantikine Human IGF-1 ELISA Kit (RSD, DG100B) based on the manufacturer’s protocol. ELISA data were analyzed using Arigo’s ELISA calculator (https://www.arigobio.com/elisa-calculator). Statistical analysis was done using GraphPad Prism 9 (GraphPad Software, San Diego, CA). A two-way repeated measures ANOVA was used to assess the effects of treatment group and time. Sidak’s multiple comparisons test was used as a post hoc test to identify significant differences between the groups at each time point. A P value < 0.05 was considered statistically significant. The data are presented as the mean ± SEM.

### Luciferase assay for estimating cancer cell proliferation

From Day 7 to Day 21, FPCO and iMac-FPCO were cultured in basic-FPCO medium. Each time point of the iMac-FPCOs was subject to the luciferase assay. iMac-FPCO and FPCO were fully lysed using a QIAGEN Tissue Lyser LT (QIAGEN) in 1 X Glo Lysis Buffer (Promega). Luciferase assays were carried out using the One-Glo Luciferase Assay System (Promega) based on the manufacturer’s protocol.

### Gemcitabine resistance assay

Luc-EGFP-PDAC cells were used to generate the FPCO and iMac-FPCO. FPCO and iMac-FPCO for the drug resistance assay were prepared on cell culture inserts placed in 12-well plates. The PCO was prepared by seeding 3 × 10^4^ PDAC cells in a 1:1 mixture of Matrigel- and DMEM-coated 96-well plates. FPCO, iMac-FPCO, and PCO were treated with gemcitabine (0, 1, and 3 μM) for 72 h. During drug treatment, the medium was replaced with fresh medium containing each drug every 24 h. After drug treatment, each organoid was collected, and the number of cancer cells was quantified using the luciferase assay.

### Macrophage depletion assay in iMac-FPCO

To enable the inducible depletion of iMacs, a stable hiPSC line expressing a dimerizer-iCasp9 ^33^ was used for macrophage differentiation. These iCasp9-iMacs were then used to construct iMac-FPCOs as described above. To induce macrophage depletion, the chemical homodimerizer B/B (Takara Bio, #635059) was added to the culture medium at a final concentration of 1 µM, beginning on Day 7 of culture. The medium was replaced every 1–2 days until the endpoint of the experiment. As a negative control, an equivalent volume of dimethyl sulfoxide (vehicle) was added to a parallel set of iMac-FPCOs. Macrophage-free FPCOs were cultured simultaneously as a comparator group. The effect of macrophage depletion on cancer cell proliferation was evaluated on Day 21 using a luciferase assay.

### Inhibitor screening test

Luc-EGFP-PDAC cells were used to generate iMac-FPCO. For 7 days from Day14 to Day21, iMac-FPCO was incubated with Picropodophyllin (S7668, Selleck, 2 μM), Linsitinib (S1091, Selleck, 2 μM), ISO1 (S7732, Selleck, 10 μM), LSKL (P1164, Selleck, 1 μM), Vixarelimab (A3012, Selleck, 10 μM), SSR128129E (S7167, Selleck, 1 μM), ITD-1 (S6713, Selleck, 3 μM), Diacerein (S4267, Selleck, 10 μM), Crizotinib(S1068, Selleck, 1 μM), Latozinemab (A2920, Selleck, 1 μM), or Anti-TNFSF12/TWEAK (A2318, Selleck, 1 μM). During treatment, the medium was replaced with fresh medium containing each compound every 24–48 h. Each organoid was collected, and the number of cancer cells was quantified using the luciferase assay as described previously.

### scRNA sequencing for iMac-FPCO

For scRNAseq analysis, iMac-FPCO was digested into a single-cell suspension with collagenase (Sigma-Aldrich) and Pronase (Roche) as described previously.^24,25^ The library was constructed using the Chromium NEXT GEM Single Cell 3’ Kit v3.1 (PN-1000130, 10x Genomics), Chromium NEXT GEM Chip G Single Cell Kit (PN-1000127), and Chromium NEXT GEM Single Cell 3’ v3.1 Gel Beads (PN-1000129, 10x Genomics). D5000 ScreenTape (Agilent Technology) was used as a high-sensitivity quality control, and sequencing was conducted on a NovaSeq6000 (Illumina) with paired-end 150 bp reads. The raw data were processed using Cell Ranger (10x Genomics, v7.0.1). scRNA-seq data were analyzed using Seurat v5.2.1 in R. Quality control filtering retained cells with >500 genes, >1500 UMI counts, and <15% mitochondrial gene expression. Doublets were removed using scDblFinder.^51^ The data were normalized using LogNormalize, and the top 2,000 variable genes were identified. Batch correction was done using Harmony integration. UMAP visualization and cluster analysis were performed using the first 30 principal components with a resolution of 0.3. Macrophage subsets were further analyzed at a resolution of 0.5. Differential gene expression analysis was conducted using FindMarkers with log2FC ≥ 1 and adjusted p-value < 0.01.

### Patient scRNA-seq data analysis

For comparative analysis, publicly available scRNA-seq data from pancreatic cancer patients^26^ were obtained from multiple studies and integrated using Seurat v5.2.1. Myeloid cells and macrophages were extracted based on their original cell type annotations provided in the published datasets. To manage the batch effect among patients, integration was performed using Reciprocal Principal Component Analysis (RPCA). Briefly, patients with over 100 cells were selected for precise anchor detection in RPCA. Each sample was first normalized using LogNormalize, and the top 2,000 variable genes were identified, followed by scaling and principal component analysis (PCA) of these features. Batch correction was performed using RPCA, with the pre-computed PCA as the input reduction and a k-weight of 50. All downstream analyses were performed on the resulting integrated reduction. UMAP visualization and clustering analysis were conducted using the first 30 principal components and a resolution of 0.3. Macrophage subtypes were classified based on signature genes reported in the literature,^16^ including IL1B TAM, SPP1 TAM and FOLR2 TAM. Differential gene expression analysis was performed using FindMarkers with the criteria of log2FC ≥ 1 and adjusted p-value < 0.01.

### scRNA-seq reference mapping and annotation

To computationally compare the iMac-FPCO TAMs (query) with the PDAC-TAM atlas of the patient (reference), we used the reference-based mapping workflow in the Seurat R package.^29^ This approach was designed to align datasets processed with different integration techniques. The dataset for iMac-FPCO TAMs was first normalized using standard log-normalization. The patient atlas was used as a fixed reference, with pre-existing 30-dimensional RPCA reduction. We identified 2,000 shared anchor features among the datasets. Anchors were then identified by projecting the query data into the established RPCA space of the reference. Using these anchors, cell type labels from the reference metadata were probabilistically transferred to each query cell. This process generated a predicted cell type label and a quantitative mapping confidence score (from 0 to 1) for each query cell. Finally, to visually validate the mapping, the query cells were projected directly onto pre-calculated UMAP coordinates of the reference using the same anchor set. The resulting cellular composition, mapping confidence scores, and UMAP projections were used to assess the transcriptional fidelity of the organoid model against the patient atlas.

### RNA velocity analysis and fate probability analyses

RNA velocity analysis was performed using scVelo (v0.3.3)^30^. Spliced and unspliced transcript count matrices from iMac-FPCO samples on Day 7 and Day 14 were concatenated to generate a unified dataset containing 9,534 cells and 36,601 genes. RNA velocity vectors were estimated using the dynamical model. The analysis was focused on macrophage populations across four clusters. Latent time was computed to infer developmental trajectories.

To quantitatively model the differentiation trajectories and compute fate probabilities, we used CellRank (v.2.0.8).^31,32^ A cell-cell transition matrix was calculated by combining transcriptomic similarity with the RNA velocities calculated by scVelo. Using this matrix, CellRank identified terminal states of the differentiation process. Based on the data, the FOLR2 TRM-TAM 2 and IL1β TAM clusters were considered as terminal states. Next, fate probabilities towards these two terminal lineages were analyzed for all macrophages. This analysis was used to formally establish the FOLR2 TRM-TAM 1 cluster as a common progenitor by quantifying its high probability of differentiating towards both terminal fates.

### Cell-cell communication analysis

Cell-cell communication analysis was conducted using CellChat v1.6.1 in R.^34^ For iMac-FPCO co-cultures, macrophages, tumor cells, CAFs, and ECs were merged into a single Seurat object and normalized using LogNormalize. The CellChatDB human database was used, which focused on “Secreted Signaling,” “Cell-Cell Contact,” and “ECM-Receptor” interactions. Overexpressed genes and their interactions were identified, followed by communication probability calculation using the triMean method. Communications were filtered to include only those with a minimum of ten cells. Communication probability and pathway analyses were performed using computeCommunProb and computeCommunProbPathway functions. For comparative analyses, the same workflow was applied to FPCO-only cultures (tumor cells, CAFs, ECs) and patient pancreatic cancer samples^16^ (TAMs, tumor cells, CAF, EC) from independent datasets. Network centrality analyses was conducted to identify major signaling roles, and communication patterns were identified using non-negative matrix factorization with k=5 patterns for both outgoing and incoming signals.

### Statistical Analysis

Experiments were repeated at least three times. Statistical analysis was conducted using GraphPad Prism 9 (GraphPad Software). For multiple group comparisons, significance was assessed via a one-way analysis of variance followed by Tukey’s multiple comparison test for all pairwise comparisons or Dunnett’s multiple comparisons test for comparisons against the control. For comparisons involving only two groups, an unpaired Student’s t-test was used. The data are presented as mean ± standard error of the mean. In all cases, p < 0.05 was considered statistically significant.

## Data availability statement

The raw single-cell RNA sequencing data generated in this study have been deposited in the Gene Expression Omnibus (GEO) under accession number GSE311616. The data are currently private and can be accessed by reviewers using the following secure token: ubsfioeednolhgv. Any additional information required to reanalyze the data reported in this paper is available from the corresponding author upon request.

## Acknowledgements

We would like to thank all team members for providing technical support. This work was supported by the Japan Agency for Medical Research and Development AMED; JP23fk0210129, JP23bm1223007, JP23bm1523004, JP23bm1523006, JP24bk0104174 JP23ym0126126, JP24ym0126803, JP25ym0126805 to H.T., JP23gm6210029 to N.T. and the Ministry of Education, Culture, Sports, Science, and Technology (MEXT; 24H00641 to H.T.; 23K27658, 23K18572 to N.T.). We would also like to thank Enago (www.enago.jp) for reviewing the language of the manuscript. The Figures (2C, 3A, 5F, 6A, 7, S5C, and graphical abstract) were created with BioRender.com (www.biorender.com).

## Author contributions

Y.Y., K.T., S.T., N.T. and H.T., conception of the study; Y.Y., K.T., and S.T., data collection; Y.Y., K.T., and S.T., data analysis; Y.Y., K.T., S.T., and R.E., sample preparation; K.T., S.T., A.O., K.A., T.K. and N.T., data discussion; Y.Y., K.T., A.O., K.A. and N.T., manuscript writing; T.K., N.Y., Y.M., M.O., N.T. and H.T., supervision

## Declaration of Generative AI and AI-assisted technologies in the writing process

During the preparation of this work the author used Gemini for English proofreading and data analysis. After using this tool, the author reviewed and edited the content as needed and takes full responsibility for the content of the publication.

## Supplementary figure legends

**Figure S1. Establishment and validation of the TRM ratio cutoff for patient stratification, Related to Figure 1**

(A) Histograms showing the distribution of the TRM ratio (percentage of FOLR2^+^CD68^+^ cells among CD68^+^ cells) in the clinical cohort of 74 PDAC patients analyzed in this study. The red dashed line indicates the average value, whereas the blue solid line indicates the cutoff value of 10%.

(B) Histograms showing the distribution of the FOLR2-TRM ratio in a public scRNA-seq dataset. The red dashed line indicates the average value, whereas the blue solid line indicates the cutoff value of 10%.

(C) Receiver operating characteristic curve analysis for predicting 1-year overall survival using the TRM ratio as a continuous variable. The analysis identified an optimal cutoff value of 9.6%, which supports the use of a 10% threshold for patient stratification. The area under the curve and p-value are shown.

scRNA-seq, single cell RNA sequencing; TRM, tissue-resident macrophage.

**Figure S2. Gating strategy for the characterization of iMacs, Related to Figure 2**

(A) Representative flow cytometry gating strategy used to characterize and confirm the purity of iMacs on Day 21 of differentiation. Sequential gates were applied to identify single cells and live cells, followed by analysis of the expression of CD14 and CD45 expression to identify the mature macrophage population (CD14^+^CD45^+^). hiPSC, human induced pluripotent stem cell; iMac, hiPSC-derived macrophage.

**Figure S3. Additional scRNA-seq analyses comparing iMac-FPCO macrophages to patient and BMDM-like models, Related to Figure 3**

(A) Heatmap showing the expression of marker genes within cancer cells, CAFs, TAMs, and ECs in iMac-FPCO.

(B) Heatmap showing signature scores for the marker genes used to define the seven distinct TAM clusters within the patient PDAC-TAM atlas

(C) Heatmap showing the expression of key marker genes used to define the 7 distinct TAM clusters within the patient PDAC-TAM atlas.

(D) Histogram showing the maximum mapping confidence score in iMac-FPCO TAMs. The red dashed line indicates a mapping confidence score of 0.5.

(E) Integrated UMAP plot showing the distinct clusters between macrophage populations from iMac-FPCO (red) and M0-FPCO (green).

(E) Feature plot showing FOLR2-TAM signature scores on iMac-FPCO and M0-FPCO, indicating enrichment of these signatures in the iMac-FPCO TAMs.

BMDM, bone marrow-derived macrophage; scRNA-seq, single cell RNA sequencing; TAM, tumor-associated macrophage; FPCO, fused pancreatic cancer organoid; iMac-FPCO, FPCO with iMacs; iMac, hiPSC-derived macrophage; hiPSC, human induced pluripotent stem cell; PDAC, pancreatic ductal adenocarcinoma; UMAP, Uniform Manifold Approximation and Projection; EC, endothelial cell.

**Figure S4. Further characterization of macrophage subpopulations and trajectory in iMac-FPCO, Related to Figure 4**

(A) UMAP plots showing the signature scores for various patient-derived TAM gene sets (e.g., SPP1 TAMs, HSP TAMs, MT TAMs) projected onto the iMac-FPCO macrophage clusters.

(B) Volcano plots showing the differentially expressed genes comparing the FOLR2 TRM-TAM 1 cluster (left) and the FOLR2 TRM-TAM 2 cluster (right). Upregulated genes related to antigen presentation (e.g., *HLA-DPA1*) and tissue residency (e.g., *LYVE1*) are highlighted.

(C) Bar charts showing the top enriched GO terms for each of the four identified macrophage subpopulations, which provide further detail on their distinct functional profiles.

FPCO, fused pancreatic cancer organoid; iMac-FPCO, FPCO with iMacs; iMac, hiPSC-derived macrophage; hiPSC, human induced pluripotent stem cell; UMAP, Uniform Manifold Approximation and Projection; GO, Gene Ontology.

**Figure S5. Reproducibility of functional assays across individual patient-derived organoid cases, Related to Figure 5**

(A, B) Luciferase-based proliferation curves for three independent patient-derived organoid cases: (A) Patient 2, and (B) Patient 3. Organoids were cultured either with (iMac-FPCO) or without (FPCO) iMacs 21 days. For each patient line, luminescence values were normalized to the mean of the FPCO and iMac-FPCO groups on Day 7.

(C) Schematic of the inducible iCasp9 system used for the selective, CID-induced depletion of iMacs.

(D) Representative flow cytometry plots confirming the efficient depletion of CD45^+^ macrophages in iCasp9-iMac-FPCOs following treatment with CID (iMac-depletion) compared with the vehicle control.

(E, F) Quantification of cancer cell proliferation on Day 21 in the macrophage depletion experiment for two individual patient cases: (E) Patient 2, and (F) Patient 3. For each patient line, luminescence values were normalized to the mean of the corresponding iMac-FPCO group on Day 21.

(G) Representative immunofluorescence images providing visual support for the clinical data quantified in Figure 5I. The images show staining for cancer cells (CK19, green), the proliferation marker Ki-67 (red), TRMs (FOLR2, yellow), and nuclei (DAPI, blue) in tumors from representative high TRM (top) and low TRM (bottom) patients. Scale bar, 100 µm.

(H, I) Quantification of cancer cell viability in response to increasing concentrations of gemcitabine in PCO, FPCO, and iMac-FPCO cultures for two individual patient cases: (H) Patient 2, and (I) Patient 3. For each patient line, luminescence values from five independent cultures for the PCO, FPCO, and iMac-FPCO groups were normalized to the mean of their respective 0 μM controls.

For the quantitative assays, the data are presented as the mean ± SEM from 5–6 biological replicates per biological case. Statistical significance was determined by a two-way ANOVA with Sidak’s multiple comparisons test (A, B, H, I) or a one-way ANOVA with Dunnett’s multiple comparisons test (E, F).

FPCO, fused pancreatic cancer organoid; iMac-FPCO, FPCO with iMacs; iMac, hiPSC-derived macrophage; hiPSC, human induced pluripotent stem cell; CID, chemical inducer of dimerization; iCasp9, inducible caspase-9; TRM, tissue-resident macrophage; PCO, pancreatic cancer organoid; SEM, standard error of mean; ANOVA, analysis of variance.

**Figure S6. Supporting data for mechanistic analyses of cell-cell interactions, Related to Figure 6**

(A) CellChat circle plots visualizing predicted ligand-receptor interactions from CAFs and TAMs to cancer cells in iMac-FPCO (top) and a patient sample (bottom).

(B and C) Quantification of cancer cell proliferation on Day 21 in the experiment adding IGF1R inhibitor for two individual patient cases: (B) Patient 2, and (C) Patient 3. For each patient line, luminescence values were normalized to the mean of the corresponding iMac-FPCO group at Day 21.

(D) Bar chart showing the effect of the IGF1R inhibitors, Picropodophyllin and Linsitinib, on the proliferation of cancer cells in macrophage-free FPCOs. Luminescence values were normalized to the mean of FPCO (control).

(E) Dot plot quantifying IGF1 gene expression across all TAM subsets identified by scRNA-seq in the PDAC-TAM atlas.

For all quantitative assays, the data are presented as the mean ± SEM from 5–6 biological replicates per biological case. Statistical significance was determined by a one-way ANOVA with Dunnett’s multiple comparisons test (B, C, and D).

CAF, cancer-associated fibroblast; FPCO, fused pancreatic cancer organoid; iMac-FPCO, FPCO with iMacs; iMac, hiPSC-derived macrophage; hiPSC, human induced pluripotent stem cell; TAM, tumor-associated macrophage; IGF1, insulin-like growth factor 1; SEM, standard error of mean; FPCO, fused pancreatic cancer organoid; PPP, picropodophyllin.

**Table 1. Clinicopathological characteristics of patients**

Comparison of the clinicopathological characteristics between high TRM (n = 33) and low TRM (n = 41) patient groups. Continuous variables are presented as the median (range). Statistical significance was determined using the Mann-Whitney U test for continuous variables and Fisher’s exact test for categorical variables. Abbreviations: CA19-9, carbohydrate antigen 19-9; UICC, Union for International Cancer Control; TRM, tissue-resident macrophage.

**Table 2. Top ligand-receptor interactions between cell types**

Top ligand-receptor pairs between different cell types in the iMac-FPCO model, as predicted by

CellChat analysis of the scRNA sequencing data. Interactions are ranked by their communication probability, and the primary cell source of each ligand is indicated. Abbreviations: CAFs, cancer-associated fibroblasts; TAMs, tumor-associated macrophages; TRMs, tissue-resident macrophages; FPCO, fused pancreatic cancer organoid; iMac-FPCO, FPCO with iMacs; iMac, hiPSC-derived macrophage; hiPSC, human induced pluripotent stem cell; scRNA-seq, single cell RNA sequencing.

**Table 3. List of inhibitors used in this study**

Summary of the functional inhibitor screen performed on iMac-FPCO. Inhibitors targeting key ligand-receptor pairs predicted by CellChat analysis were added to the cultures on Day 14, and cancer cell proliferation was measured by luciferase assay on Day 21. The “Observed Proliferation Inhibition Rate” was calculated from the mean difference in relative luminescence between the inhibitor-treated group and the vehicle control. Statistical significance was determined by a one-way ANOVA with Dunnett’s multiple comparisons test against the vehicle control. Abbreviations: ns, not significant. *p < 0.05; **p < 0.01; ***p < 0.001; FPCO, fused pancreatic cancer organoid; iMac-FPCO, FPCO with iMacs; iMac, hiPSC-derived macrophage; hiPSC, human induced pluripotent stem cell; ANOVA, analysis of variance.

**^a^No Inhibition:** These inhibitors showed a mean difference near zero or a negative value, indicating no inhibitory effect on cancer cell proliferation in this model.

**^b^Proliferation:** Anti-TWEAK treatment resulted in a statistically significant increase in cancer cell proliferation (mean difference of −0.7933), suggesting a pro-proliferative role for TWEAK signaling in this model.

---

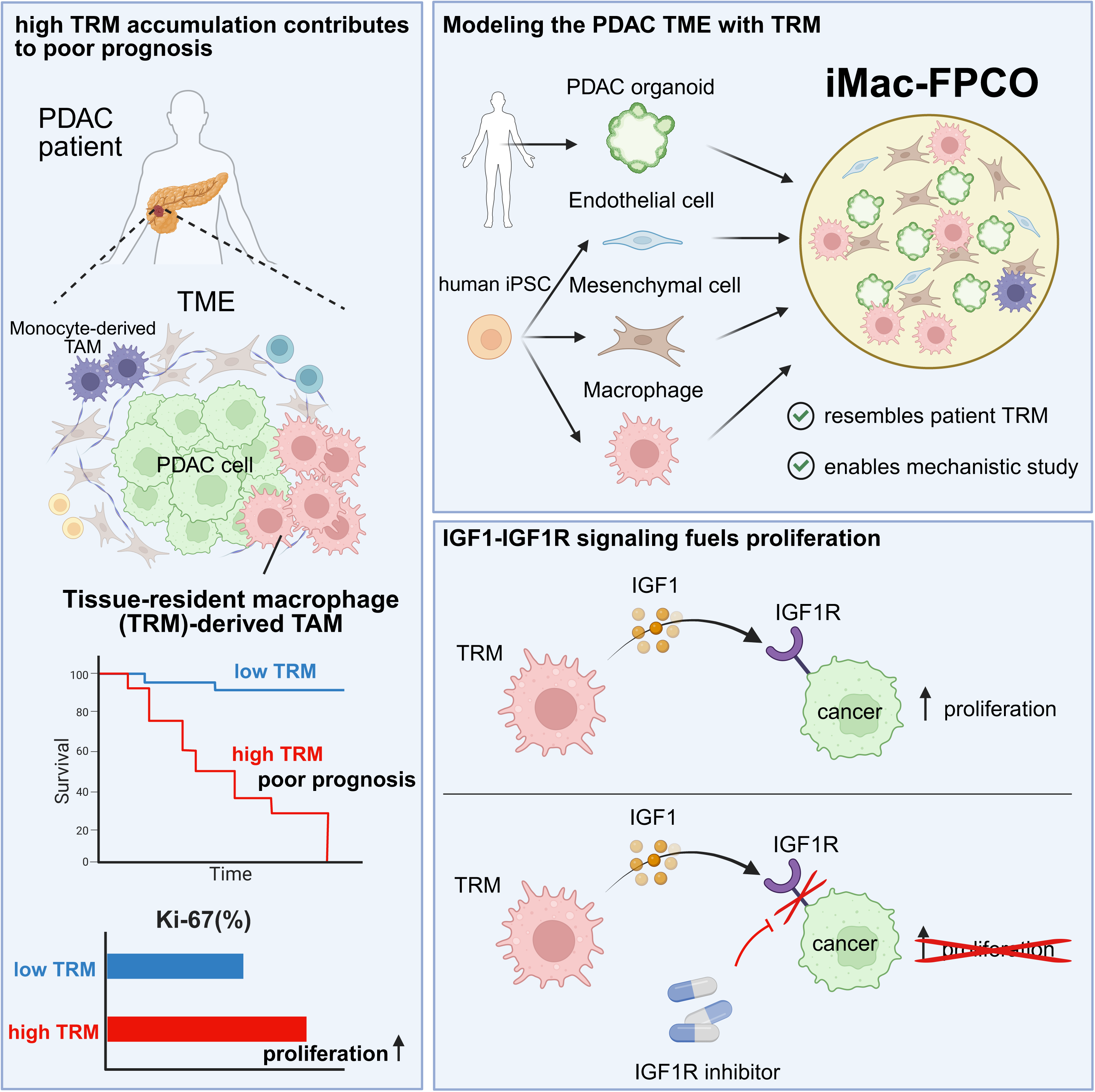

