## Supplementary figures and images for "Tumor-associated tissue-resident macrophages drive pancreatic cancer progression through IGF1-IGF1R signaling"

### Supplementary Figure

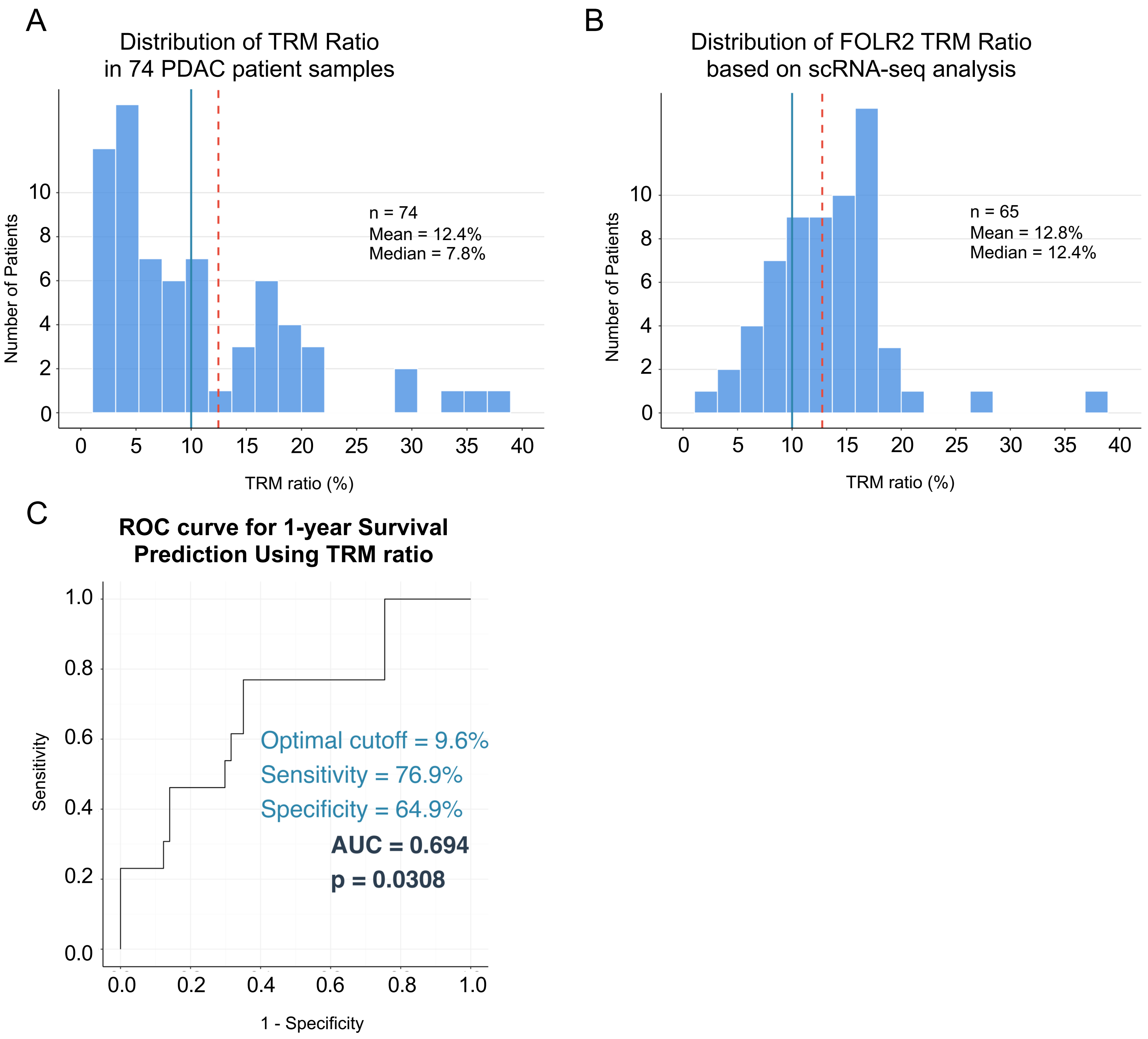

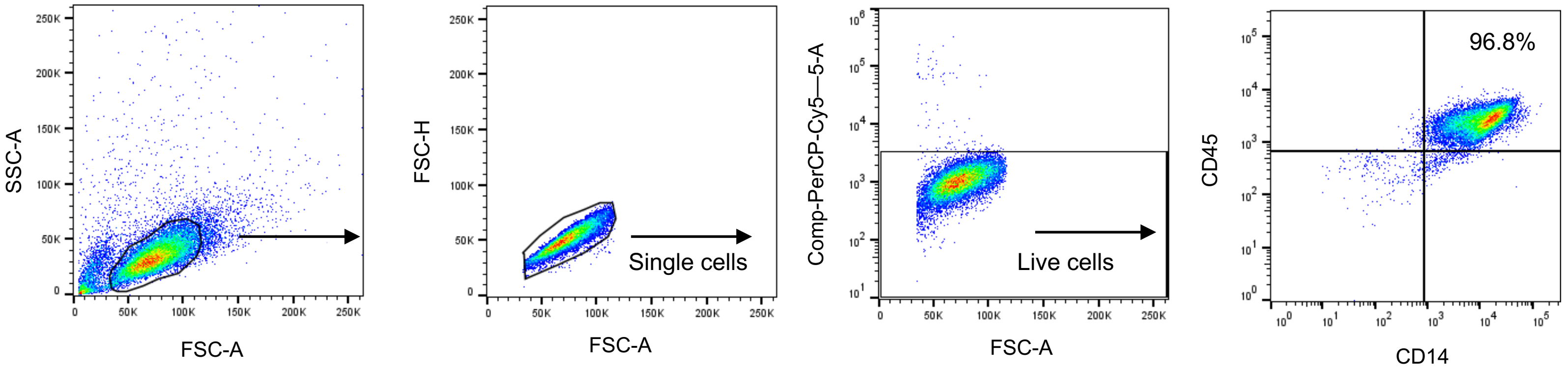

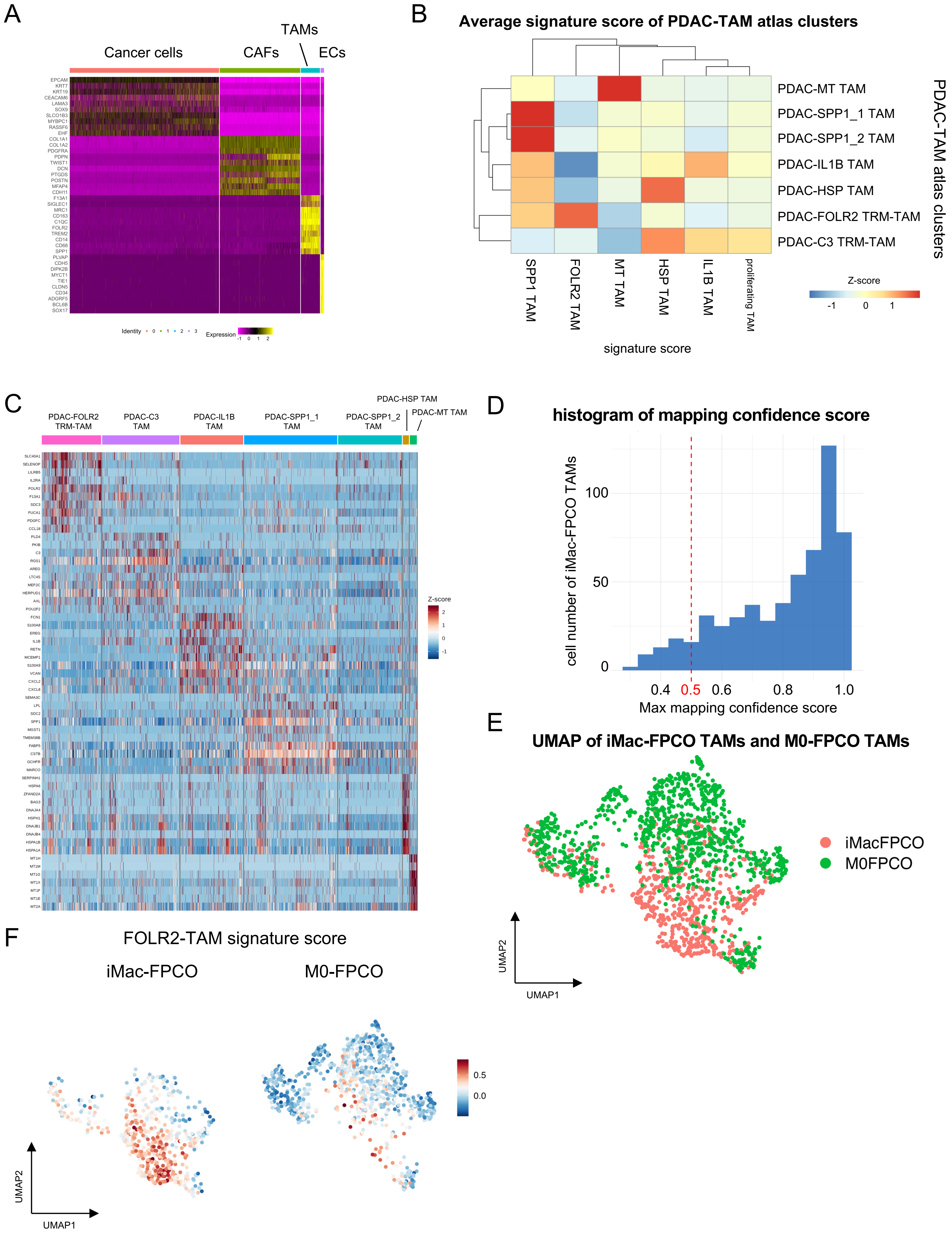

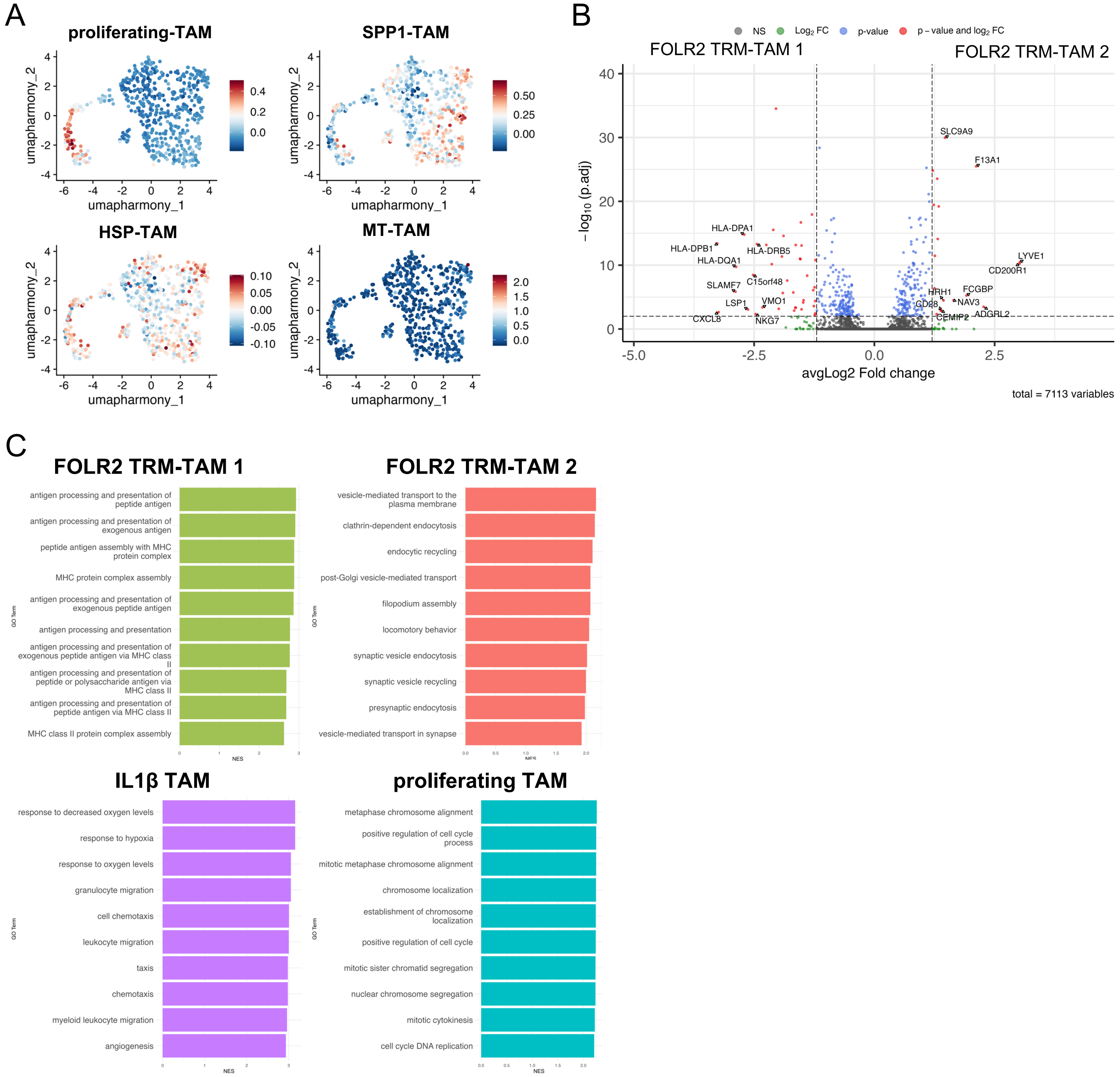

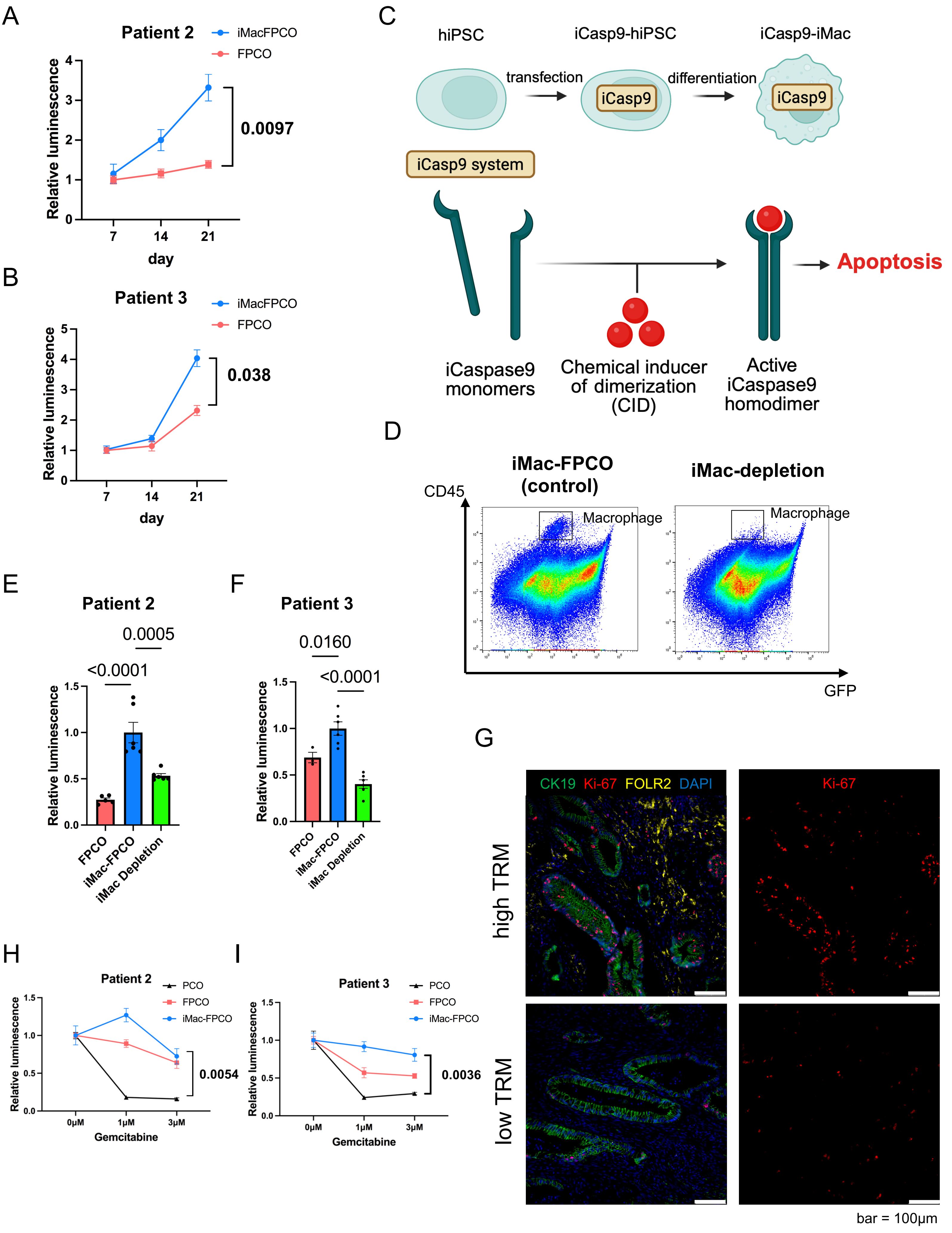

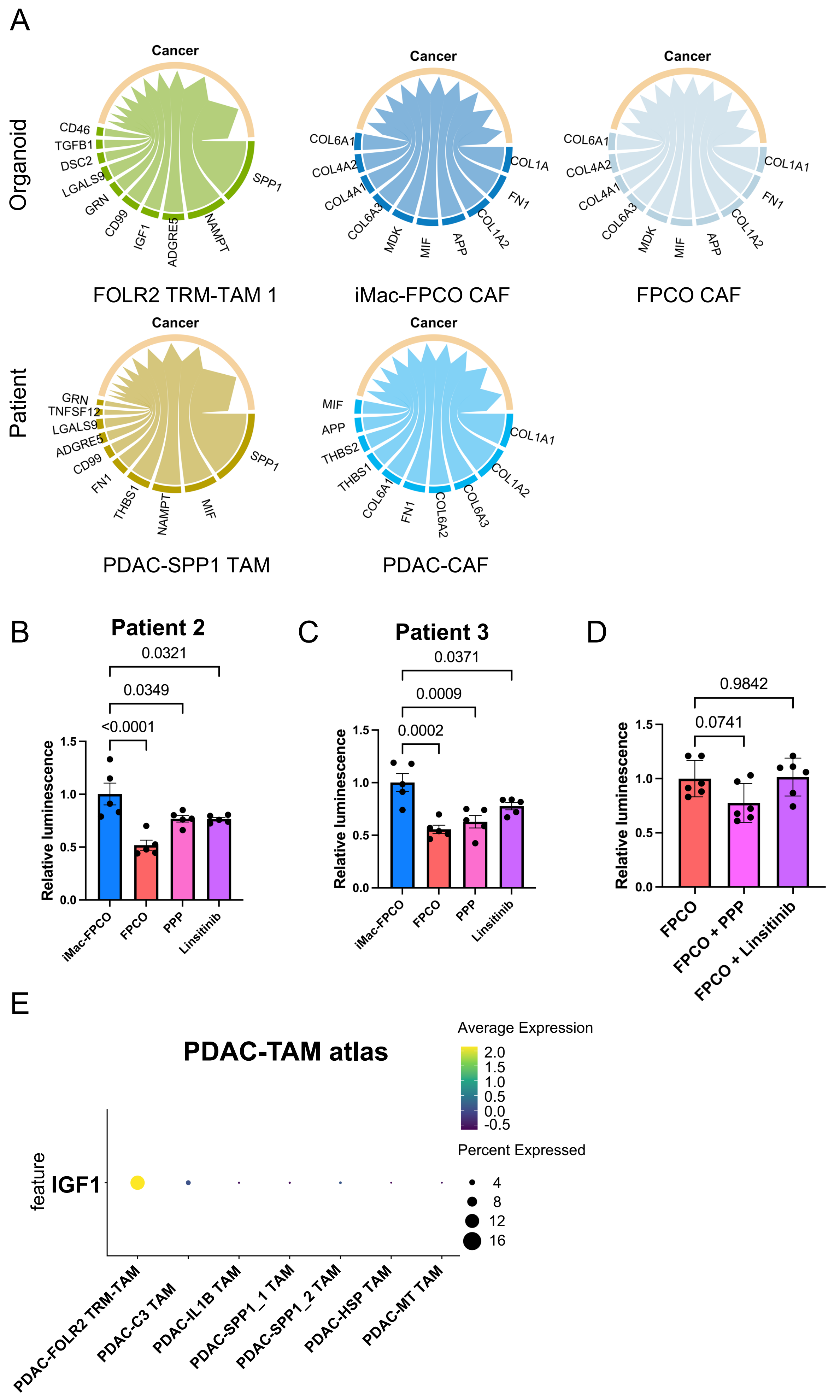
