## Supplementary material for "Tumor-associated tissue-resident macrophages drive pancreatic cancer progression through IGF1-IGF1R signaling": Table

Table 1. Clinicopathological characteristics of patients

| | high TRM (n=33) | low TRM (n=41) | P value |
|---|---|---|---|
| Sex (Male/Female) | 15/18 | 23/18 | 0.48 (ns) |
| Age (y.o. median: range) | 72 (54-82) | 70 (53-88) | 0.84 (ns) |
| Preoperative CA19-9 level (U/ml median: range) | 67.9 (0-1700) | 124.1 (0.1-5590) | 0.13 (ns) |
| Histological grade (well /mod /por) | 10/17/5 | 13/22/6 | 1.00 (ns) |
| Tumor size (mm, median: range) | 32 (12-60) | 30 (10-41) | 0.14 (ns) |
| Portal vein invasion (+/-) | 13/20 | 10/31 | 0.21 (ns) |
| Arterial invasion (+/-) | 2/31 | 4/37 | 0.69 (ns) |
| Lymphatic invasion (0,1 /2,3) | 17/16 | 28/13 | 0.16 (ns) |
| Venous invasion (0,1/2,3) | 27/6 | 34/7 | 1.00 (ns) |
| UICC-stage (I /II /III /IV) | 10/7/13/3 | 10/18/12/1 | 0.18 (ns) |
| T-stage (T1 /2 /3 /4) | 5/24/3/1 | 9/27/3/2 | 0.86 (ns) |
| N-stage (N0/1/2) | 11/8/14 | 13/17/11 | 0.23 (ns) |
| M-stage (M0/M1) | 30/3 | 40/1 | 0.32 (ns) |
| Resectability(R0, R1,2) | 25/8 | 28/13 | 0.61 (ns) |
| Local recurrence (+/-) | 11/20 | 12/29 | 0.62 (ns) |
| Hematogenous recurrence (+/-) | 12/20 | 12/29 | 0.61 (ns) |
| Lymph node recurrence (+/-) | 4/28 | 5/36 | 1.0 (ns) |
| Peritoneal dissemination recurrence (+/-) | 7/25 | 1/40 | 0.018 (*) |
| Ki-67 positive rate (%, median: range) | 5.88 (0.50–17.91) | 4.66 (0.12–9.94) | 0.025(*) |

### Table 2. Top ligand-receptor interactions between cell types

| Ligand | Receptor | Communication Probability | Source of Ligand |
|---|---|---|---|
| SPP1 | CD44 | 0.18666 | All macrophages |
| SPP1 | ITGAV_ITGB1 | 0.16901 | All macrophages |
| NAMPT | INSR | 0.12555 | All macrophages |
| MIF | CD74_CD44 | 0.12312 | IL1B TAM |
| SPP1 | ITGAV_ITGB6 | 0.11521 | All macrophages |
| SPP1 | ITGAV_ITGB5 | 0.10551 | All macrophages |
| THBS1 | CD47 | 0.06267 | IL1B TAM |
| THBS1 | SDC1 | 0.06267 | IL1B TAM |
| THBS1 | SDC4 | 0.05470 | IL1B TAM |
| PTN | NCL | 0.04551 | CAFs |
| THBS1 | ITGA3_ITGB1 | 0.04418 | IL1B TAM |
| IGF1 | IGF1R | 0.03537 | FOLR2 TRM-TAM |
| IGF1 | ITGA6_ITGB4 | 0.03197 | FOLR2 TRM-TAM |
| ADGRE5 | CD55 | 0.02936 | LYVE1(-) |
| CD99 | CD99 | 0.02438 | IL1B TAM |
| PTN | SDC1 | 0.02325 | CAFs |
| HBEGF | EGFR | 0.02042 | IL1B TAM |
| HBEGF | EGFR_ERBB2 | 0.02027 | All macrophages |
| PTN | SDC4 | 0.02018 | CAFs |
| GRN | SORT1 | 0.01987 | FOLR2 TRM-TAM |
| LGALS9 | CD44 | 0.01973 | All macrophages |
| MDK | LRP1 | 0.01939 | CAFs |
| DSC2 | DSG2 | 0.01592 | All macrophages |
| TGFB1 | ACVR1B_TGFBR2 | 0.01496 | All macrophages |
| TGFB1 | TGFBR1_R2 | 0.01082 | All macrophages |
| TNFSF12 | TNFRSF12A | 0.01020 | All macrophages |
| TGFB1 | ACVR1_TGFbR | 0.00794 | IL1B TAM |
| SEMA6B | PLXNA2 | 0.00656 | IL1B TAM |
| SEMA3C | NRP2_PLXNA2 | 0.00565 | CAFs |
| TNF | TNFRSF1A | 0.00327 | IL1B TAM |
| OSM | OSMR_IL6ST | 0.00237 | IL1B TAM |
| SEMA3C | NRP2_PLXNA3 | 0.00188 | CAFs |
| HGF | MET | 0.00165 | CAFs |
| FGF7 | FGFR1 | 0.00135 | CAFs |
| IL1B | IL1R2 | 0.00128 | IL1B TAM |
| IL1A | IL1R2 | 6.88E-05 | CAFs |

Table 3. List of inhibitors used in this study

| Inhibitor | Target | Concentration | Observed Proliferation Inhibition Rate | Adjusted P value |
|---|---|---|---|---|
| Picropodophyllin | IGF1R | 2.0µM | 60.8% | 0.0018 (**) |
| Linsitinib | IGF1R | 2.0µM | 48.1% | 0.0216 (*) |
| ISO-1 | MIF | 10.0µM | 21.70% | 0.7177 (ns) |
| LSKL | THBS1 | 1.0µM | 20.90% | 0.7526 (ns) |
| Vixarelimab | OSMR | 10.0µM | 14.60% | 0.9653 (ns) |
| Crizotinib | HGF-MET | 1.0µM | 13.00% | 0.9856 (ns) |
| SSR128129E | FGFR1 | 1.0µM | 8.60% | 0.9991 (ns) |
| ITD-1 | TGFb | 3.0µM | No Inhibition [a] | 0.9991 (ns) |
| Diacerein | IL1B | 10.0µM | No Inhibition [a] | 0.9991 (ns) |
| Latozinemab | GRN-SORT1 | 1.0µM | No Inhibition [a] | 0.7941 (ns) |
| Anti-TWEAK | TWEAK | 1.0µM | Proliferation [b] | 0.0003 (***) |
